# Hair-bearing human skin generated entirely from pluripotent stem cells

**DOI:** 10.1101/684282

**Authors:** Jiyoon Lee, Cyrus Rabbani, Hongyu Gao, Matthew Steinhart, Benjamin M. Woodruff, Zachary Pflum, Alexander Kim, Stefan Heller, Yunlong Liu, Taha Z. Shipchandler, Karl R. Koehler

## Abstract

The skin is important for regulating bodily fluid retention and temperature, guarding against external stresses, and mediating touch and pain sensation. The skin is also susceptible to damage from burns, diseases, or genetic defects, which affect nearly one billion people worldwide^1,2^. For the advancement of skin regenerative therapies, it remains challenging to construct new skin with hair follicles and nerves in tissue cultures and in bioengineered skin grafts^3–8^. Here, we report an organoid culture system that generates complex skin from human pluripotent stem cells. We use step-wise modulation of the TGFβ and FGF signalling pathways to co-induce cranial epithelial cells and neural crest cells within a spherical cell aggregate. During 4-5 months incubation, we observe the emergence of a cyst-like skin organoid composed of stratified epidermis, fat-rich dermis, and pigmented hair follicles equipped with sebaceous glands. A network of sensory neurons and Schwann cells form nerve-like bundles that target Merkel cells in organoid hair follicles, mimicking human touch circuitry. Single-cell RNA sequencing and direct comparison to foetal specimens suggest that skin organoids are equivalent to human facial skin in the second-trimester of development. Moreover, we show that skin organoids produce planar hair-bearing skin when grafted on nude mice. Together, our results demonstrate the self-assembly of nearly complete skin tissue *in vitro* that can be used to reconstitute skin *in vivo*. We anticipate that our skin organoid model will be foundational to future studies of human skin development, disease modelling, or reconstructive surgery.

---

For decades, culture systems containing epidermal and dermal cells have been used to model human skin outside of the body^3,9,10^; however, it has been persistently difficult to grow and maintain functional skin appendages in culture. We recently reported that appendage-bearing skin organoids can be generated via directed differentiation of epidermal and dermal cells from mouse pluripotent stem cells (PSCs)^4^. From this study, we recognised that a 3D cyst composed of surface ectoderm (i.e. epidermal precursors) enveloped by mesenchymal cells (i.e. dermal precursors) creates a self-organizing system that mimics the cell-to-cell signalling mechanisms needed for embryonic hair follicle (HF) induction. Thus, to produce a humanized skin organoid model, we sought to co-induce surface ectodermal and mesenchymal cells from human PSCs (hPSCs) to mimic normal skin development (Fig. 1a, Extended Data Fig. 1). As a starting point, we revisited our group’s human inner ear organoid induction method^11^, which yields epidermal keratinocytes and cranial neural crest cell (CNCC)-derived mesenchymal tissues as by-products. Using a modified approach, we dissociated hPSCs into single cells and plated them in U-bottom 96- well plates to create uniform cell aggregates (Fig. 1b). We then transferred the aggregates to new plates containing a differentiation medium with key factors to promote epidermal induction: Matrigel, bone morphogenetic protein-4 (BMP), and a transforming growth factor beta (TGFβ) inhibitor, SB431542 (SB; see **Methods** and **Supplementary Note 1** for protocol details). TGFβ inhibition has been shown to promote ectoderm induction from pluripotent stem cells, whereas BMP activation promotes surface ectoderm (also known as non-neural ectoderm) induction and suppresses neural induction^12–15^. This differentiation strategy produced uniform epithelial cysts, ∼500-1000 µm in diameter, composed of TFAP2A^+^ ECAD^+^ epithelial cells by day 30 (Fig. 1c; upper row, Extended Data Fig. 2). These results were replicable using a *Desmoplakin (DSP)-GFP* human induced-PSC (hiPSC) line, in which desmosomes at the apical surface of the nascent epithelium are *GFP^+^* (Fig. 1d, **Supplementary Fig. 1**). However, over long-term culture (>60 days), epidermal cysts rarely displayed signs of higher-order skin morphology, such as epithelia with stratified layers of keratinocytes. Therefore, we sought additional stimuli that would induce dermal tissue to promote epidermal-mesenchymal crosstalk.

**Figure 1.**
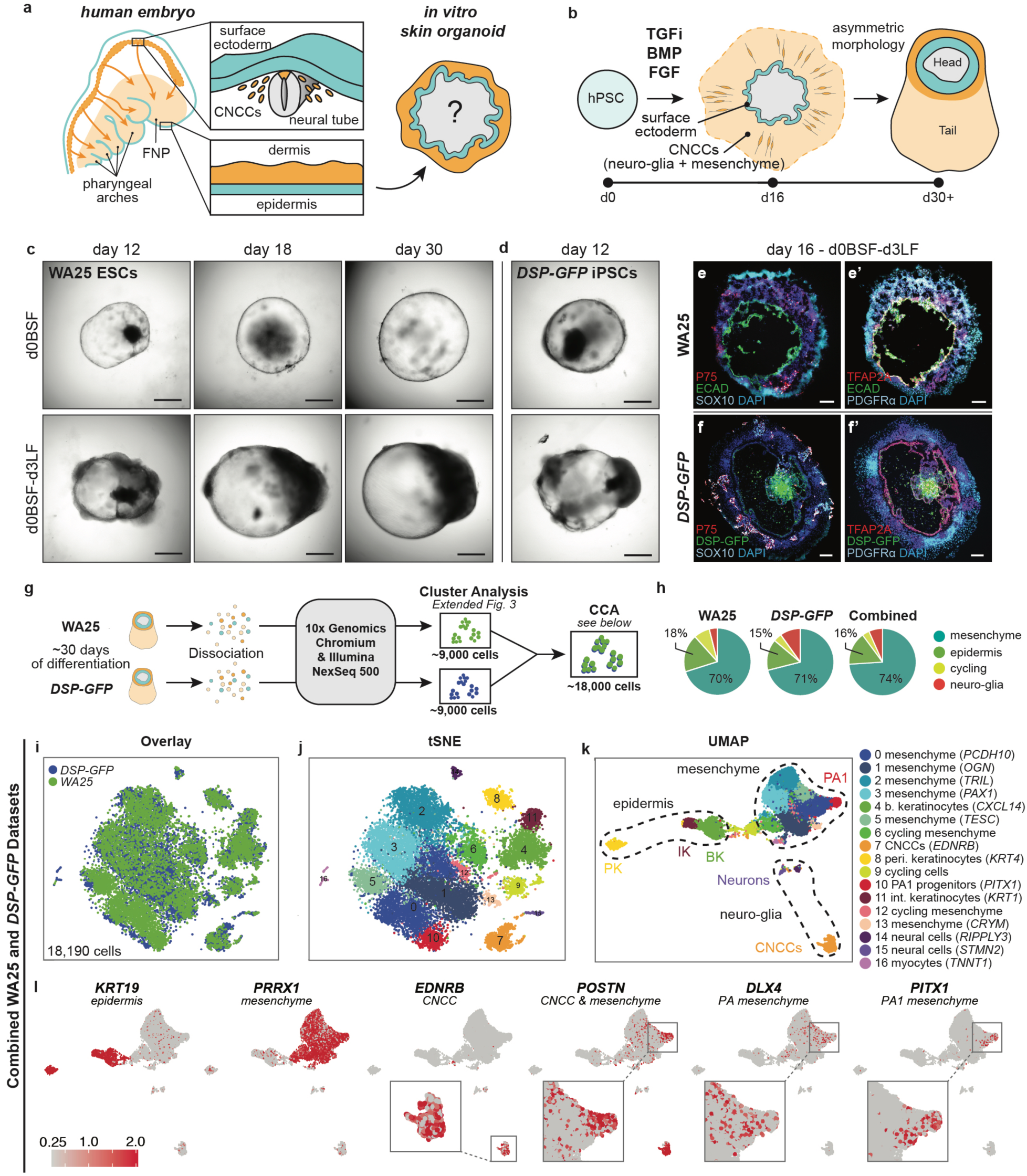
Surface ectoderm and CNCC co-induction from ESCs and iPSCs. **a**, Schematic overview of the study. **b**, Overview of skin organoid induction protocol highlighting the key timepoints and signalling pathways that are modulated. Cranial neural crest cells, CNCCs. **c, d**, Brightfield images of (c) WA25 aggregates on days 12, 18, and 30, and (d) *DSP-GFP* aggregates on day 12, comparing morphologies between d0BSF (*upper*) vs. d0BSF-d3LF treatments (*lower*). **e-f’** Immunostaining images of day-16 (e, e’) WA25 and (f, f’) *DSP-GFP* aggregates with co-induced epithelium and CNCCs under optimized d0BSF-d3LF treatment culture. (e, f) P75 and SOX10 highlight neuro-glial-associated CNCCs, and (e’, f’) PDGFRα and TFAP2A highlight mesenchyme-associated CNCCs. Epithelium is highlighted by ECAD and TFAP2A double-positive signals. **g**, Schematic of scRNA-seq experimental design. **h**, Pie charts showing the distribution of mesenchymal, epidermal, neuro-glial, and actively cycling cells across both datasets and the combined dataset. **i**, Canonical correlation analysis (CCA) plot comparing clusters of *DSP-GFP* and WA25 cells—18,190 total cells. **j**, tSNE representation of the combined cell clusters—16 unique cell subtypes were identified. **k**, UMAP representation of the combined cell clusters. The presumptive cell identities, based on *a priori* knowledge of marker genes, are listed to the right. Abbreviations (Abbr): basal (b) keratinocytes (k); cranial neural crest cells (CNCCs); peridermal (p, peri) keratinocytes; pharyngeal arch (PA)-1 progenitors; intermediate (i, int) keratinocytes. **l**, Key gene expression markers for cell subtypes and pharyngeal arch-1 identity. Scale bars, 500 µm (**c, d**), 100 µm (**e-f’**). Additional data in Extended Data Figs. 1-3.

In the embryo, dermal fibroblasts have two origins: mesoderm lineage cells (body skin and the scalp) and CNCCs (facial skin)^16,17^. In our inner ear culture system, we determined that a co-treatment of basic-FGF (FGF-2) and a BMP inhibitor, LDN-193189 (LDN) promotes induction of CNCC-like cells that can generate mesenchymal cells; however, we have not previously investigated dermal cell specification in this system^11^. Here, we optimized the timing of treatment to day-3 of differentiation, which reproducibly generated epithelial cysts that were fully covered by CNCC-derived cells (Fig. 1c, d; **Supplementary Fig. 1**, **Supplementary Table 1b, c**). To mature the organoids, we incubated them with constant agitation on an orbital shaker. We observed cells migrating radially from the epithelium between days 16-30 of differentiation (Fig. 1e, f, Extended Data Fig. 2). Both WA25 and *DSP-GFP* organoids gradually became bipolar, with the epidermal cyst partitioned to one pole (referred to as the *head*) and a mass of CNCC derivatives on the opposite pole (referred to as the *tail*; Fig. 1c, d, **Supplementary Fig. 1**). Based on immunostaining at day 16, the FGF/LDN-treated organoids consisted of an inner mantle of TFAP2A^+^ ECAD^+^ epithelial cells surrounded by an outer layer of migratory TFAP2A^+^ CNCC-like cells. Within the migratory population, there appeared to be two types of CNCC-like cells expressing either the mesenchyme-associated marker PDGFRα or the neuro-glial-associated markers SOX10 and P75 (Fig. 1e, f, Extended Data Fig. 2)^18^.

Next, we used single-cell RNA sequencing (scRNA-seq) to classify organoid cell types after one month of differentiation (29 days). Datasets from both WA25 and *DSP-GFP* organoids were analysed individually and then combined using canonical correlation analysis (CCA) to identify differences and similarities in cellular composition between cell lines (Fig. 1g). Unbiased clustering using Seurat 2 generated 15 clusters from 9,268 WA25 cells and 19 clusters from 9,013 *DSP-GFP* cells. We manually partitioned these clusters into four major cell groups: mesenchymal cells (70-71%), epidermal cells (15-18%), neuro-glial cells (4- 10%), and actively cycling *CENPF^+^* cells (4-8%) (Fig. 1h, **Supplementary Data 1, 2**). The combined CCA dataset (18,190 cells) had 16 consensus cell clusters (Fig. 1i-k, Extended Data Fig. 3**, Supplementary Data 3**). The epidermal group contained three distinct keratinocytes: *CXCL14^+^* basal, *KRT1^+^* suprabasal (also known as *intermediate*), and *KRT4^+^* peridermal keratinocytes (Fig. 1k)^19^. The mesenchymal group contained ten distinct populations, including two dermal fibroblast-like subtypes expressing *TRIL*, *TWIST2*, *ACPDD1*, and *PAX1* (Extended Data Fig. 3a-c, e-g)^20^. There was a cluster of CNCC-like cells expressing markers, such as *S100β*, *POSTN*, *EDNRB*, *CDH6*, *PLP1*, and *SOX10* (Fig. 1k, l, **Supplementary Data 3**). Two neural subgroups contained cells that express either *SOX2* (likely neural progenitors) or *STMN2*, *NEFM*, and *ELAVL2* (likely immature neurons) (Fig. 1k, **Supplementary Data 1-3**)^21^. Interestingly, a population of myelin protein zero (*MPZ*)- positive Schwann-like cells appeared in the *DSP-GFP* dataset at this stage (Cluster 14 in Extended Data 3e, f, **Supplementary Data 2**). Moreover, *TNNT1*^+^ *MYOG^+^* myocyte-like cells were more abundant in *DSP-GFP* vs. WA25 organoids (∼39:1 ratio, Cluster 16 in Extended Data 3e, f, **Supplementary Fig. 2**, **Supplementary Data 1-3**). Despite these differences, our data suggest that the overall cellular composition of organoids is remarkably similar between these two cell lines. Differences may be attributable to limited biological replicates or differential expression of imprinted genes, such as *MEG3* and *PEG3*, which were expressed in WA25, but not in *DSP-GFP* organoids (**Supplementary Fig. 2, Supplementary Data 1-3**)^22^. Notably, we did not detect progenitors for endothelial or immune cells, such as Langerhans cells and macrophages, likely due to the absence of mesoderm lineage cells in skin organoids.

During vertebrate cranial development, the pharyngeal arches (PAs) and the frontonasal process (FNP) fill with CNCC-derived mesenchymal progenitor cells migrating from the neural tube^23^. Various gene markers that pinpoint anatomical positioning among CNCC progenitors have been identified^16^. Thus, we inspected our scRNA-seq datasets for regional gene markers. Both *DSP-GFP* and WA25 organoids contained a population of *PRXX1^+^DLX1/4/5^+^* cells, reminiscent of PA or FNP mesenchymal progenitor cells (Fig. 1l, Extended Data Fig. 3d, h)^16^. Interestingly, the subset of *PRXX1^+^DLX1/4/5^+^* mesenchymal cells also expressed *PITX1*, which is a marker of aboral PA1 mesenchymal progenitor cells (Fig. 1l)^24^. Furthermore, we noted a lack of *HOXA2* (or any *HOX* gene) expression, which would indicate more caudal PAs (i.e. PA2-4) (**Supplementary Fig. 3**)^25^. Together, these data suggest that skin organoids may represent foetal skin associated with PA1 derivatives, such as the chin, cheek, and auricle skin.

After ∼50 days of differentiation, skin organoid epithelia were clearly stratified into basal, intermediate, and periderm layers (Fig. 2a). We performed additional scRNA-seq on day-48 WA25 organoids; however, for this experiment we dissected off the tail region to bias the dataset toward skin-related subtypes. As before, cells were clustered into mesenchymal, epidermal, neuro-glial, and cycling cells (Fig. 2b, c, Extended Data Fig. 4a-d, **Supplementary Data 4**). In these data, we saw more definitive markers of the dermal lineage compared to day-29. In particular, five *VIM^+^* clusters (2-6) contained cells expressing *DPT, TWIST2,* and *LUM,* as well as numerous collagens, such as *COL1A1/2*, *COL3A1*, *COL5A3*, *COL6A3, COL12A1,* and *COL21A1* (**Supplementary Data 4**)^20^. We also gained insight into self-organizing signalling mechanisms that may be governing interactions between the epidermal and dermal layers. Genes encoding WNT signalling pathway modulators were differentially expressed between epidermal (*WNT6, LEF1*) and dermal (*SFRP2*, *TCF4*, *WIF1*, *APCDD1*) cell populations (Fig. 2c, Extended Data Fig. 4e, f)^26^. Presumptive dermal cells in Cluster 2, 3, 4, and 6 expressed *FGF7* (also known as Keratinocyte Growth Factor, *KGF*), which may be a key driver of epidermal stratification in this system (Extended Data Fig. 4g)^27,28^. In addition to these findings, we identified a small subset of basal keratinocyte-like cells that expressed genes associated with Merkel cell identity, such as *ATOH1*, *KRT8/18/20*, *SOX2*, and *ISL1* (Cluster 11; Fig. 2b, d, Extended Data Fig. 4a, b)^29^. These data show that skin organoids generate diverse cell populations consistent with embryonic skin.

**Figure 2.**
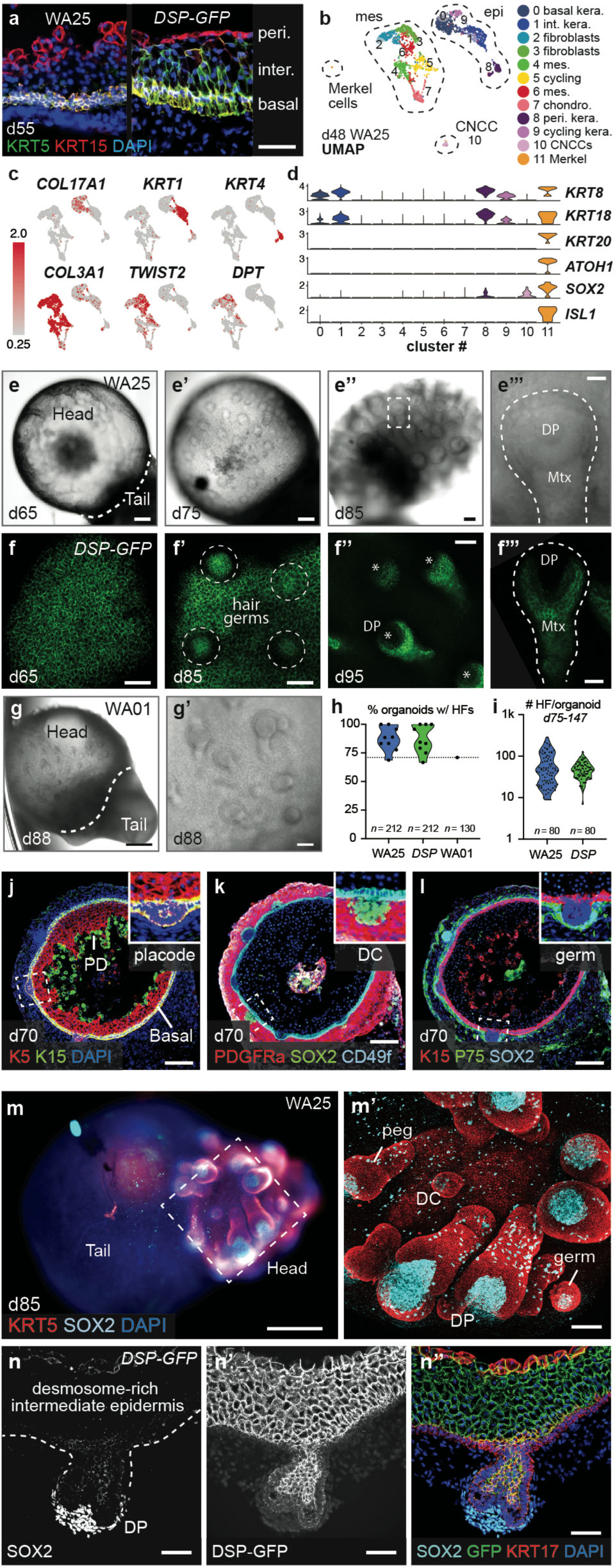
Keratinocyte and dermal differentiation mimics normal human development. **a,** Comparison of KRT5, KRT15, and TFAP2A expression in day-55 WA25 and *DSP-GFP* skin organoids. **b**, UMAP representation of scRNA-seq data from five day-48 WA25 skin organoids. Presumptive cell cluster identities were selected based on a *priori* knowledge of marker genes. Abbr: keratinocytes (kera); intermediate (int, inter); mesenchyme (mes); chondrocytes (chondro); peridermal (peri); cranial neural crest cell (CNCC). **c,** UMAP plots showing expression patterns for key markers for epidermal, mesenchymal, and CNCC populations. **d**, Violin plots showing expression levels for Merkel cell markers. **e-e’’’**, Brightfield images showing HF induction on the surface of a WA25 skin organoid between 65-85 days of differentiation. Dashed box (e’’) indicates magnified HF shown in e’’’, and dashed line (e’’’) outlines the HF. Abbr: dermal papilla (DP); matrix (Mtx). **f-f’’’**, Maximum intensity images of confocal z-stacks of the surface of multiple *DSP-GFP* organoids between 65-85 days of differentiation. Dashed circles (f’) indicate developing hair germs, asterisks (f’’) mark dermal papilla of individual hair pegs, and dashed line (f’’’) outlines a HF. **g, g’,** Brightfield images showing HF induction on the surface of a WA01 skin organoid on day 88 of differentiation. **h, i**, Violin plots showing (h) frequencies of HF formation in WA25 (87.4 ± 3.5%, 181 hairy-organoids, n=212, 9 experiments), *DSP-GFP* (87.2 ± 4.1%, 177 hairy-organoids, n=212, 9 experiments), and WA01 (71%, 92 hairy-organoids, n=130, 1 experiment) cultures (see **Supplementary Table 1** for details), and (i) average number of HFs formed in WA25 (average 64 HFs per organoid, the least=9 and the most=285 HFs per organoid, n=80) and *DSP-GFP* (average 48 HFs per organoid, the least=7 and the most=128 HFs per organoid, n=80) cultures between days 75-147. **j-l**, Immunostaining of a day-70 WA25 skin organoid with hair placodes. Antibodies were chosen to highlight epidermial layers (KRT5, KRT15, CD49f), peridermal layer (KRT15), dermis (PDGFRα), and dermal condensate cells (SOX2, P75). Abbr: periderm (PD); dermal condensate (DC). Dashed boxes indicate magnified regions presented on the right top corner of each image. **m, m’**, (m) Low magnification image of wholemount immunostained day-85 WA25 skin organoid with tail structure. (m, m’) KRT5 highlights the epidermis and outer root sheath of follicles. SOX2 marks dermal condensate and papilla cells, as well as Merkel cells and migratory melanocytes. See **Supplementary Video 1**. Dashed box indicates magnified area shown in m’. **n-n’’**, Immunostaining of a day-95 *DSP-GFP* skin organoid showing (n) a hair peg with SOX2^+^ dermal papilla, (n’) GFP^+^ desmosome-rich intermediate epidermis, and (n’’) KRT17^+^ periderm layer and outer root sheath of the follicle. Scale bars, 250 µm (**g**, **m**), 100 µm (**e-e’’, f’’, j-l**), 50 µm (**a**, **f’, g’, m’-n’’**), 25 µm (**e’’’, f, f’’’**). Additional data in Extended Data Figs. 4, 5, 6.

Human foetal HFs typically develops during weeks 9-10 (days 63-70) of gestation^30^; thus, we waited over 70 days for skin organoids to reach a hair-bearing stage. After a period of quiescence (days 30-65), hair germ-like buds emerged radially outward from the organoid surface (Fig. 2e, f, Extended Data Fig. 5). On average, hair germs appeared at 70 ± 5 days for WA25 (*n* = 80) and 72 ± 4 days for *DSP-GFP* (*n* = 80) organoids. Through day 120 of differentiation, we observed robust HF induction from batch-to-batch and across cell lines (87.4% WA25 with HFs, 9 experiments; 87.2% *DSP-GFP* with HFs, 9 experiments; Fig. 2h, Extended data Fig. 5a, b, **Supplementary Table 1a**). Hair-bearing skin organoid generation was not restricted to these two cell lines or the Koehler laboratory. In a protocol validation experiment, hair-bearing skin organoids were successfully generated from the WA01 (H1) hESC line in the Heller laboratory (70.8% WA01 with HFs, 1 experiment; Fig. 2g, h, Extended data Fig. 5c; see ***protocol validation*** in **METHODS**). By manual counting, we estimated an average of 64 HFs per WA25 and 48 HFs per *DSP-GFP* organoid (n=80 per group) (Fig. 2i). As in nascent mouse HFs, immunostaining revealed that epidermal germ cells were LHX2^+^ PCAD^+^ EDAR^+^ and dermal condensates were SOX2^+^ P75^+^ (Fig. 2j-n, Extended Data Fig. 6). The transcription factor SOX2 was expressed in every dermal papilla (DP) cell population examined (Fig. 2k-n, **Supplementary Video 1, 2**), consistent with human foetal HFs and guard, awl, and auchene mouse HFs^31^. These data show that skin organoid HFs undergo morphological transformations similar to well-documented stages—stage-1 (placode), -2 (germ), and -3 (peg), etc.—of mammalian HF induction^32^. Moreover, organoid HFs are periodically spaced throughout the epidermis, which suggests that patterning mechanisms seen in vertebrate embryos are preserved in the organoid model^33,34^.

After >100 days in culture, skin organoids had the morphology illustrated in Fig. 3a. The external appearance of skin organoids was comparable to day-132 human foetal skin viewed from the dermal-side (Fig. 3b, c). Using electron microscopy, we found the organoid HFs had all of the unique cellular layers associated with mammalian hair (Fig. 3d). Remarkably, organoid HFs were often pigmented, which led us to closely examine melanocyte development. 53.5% of WA25 organoids (n = 137, 8 experiments) and 76.2% of *DSP-GFP* organoids (n = 80, 5 experiments) had pigmented HFs (Fig. 3e). In pigmented skin organoids, we found premelanosome protein (PMEL)^+^ melanocytes evenly distributed throughout the epithelium and concentrated in the matrix region of HFs (Fig. 3f-j, Extended Data Fig. 7**, Supplementary Video 3**). Electron microscopy confirmed production of melanosomes in melanocytes of the epithelium and HFs (Fig. 3h, Extended Data Fig. 7a). Another interesting feature of late-stage organoids was hyaline cartilage, which developed in the tail region (Fig. 3a, b, Extended Data Fig. 8). *COL9A1-3^+^* chondral progenitors were evident in our day-48 scRNA-seq dataset (Extended Data Fig. 8a). Additionally, lipid-rich adipocytes developed around organoid HFs, mimicking the development of hypodermal fat (Fig. 3b, c, Extended Data Fig. 9a-d)^17^. These data reinforce a tissue architecture model whereby skin organoids represent craniofacial epidermal, dermal, and chondral tissues developing *inside-out*.

**Figure 3.**
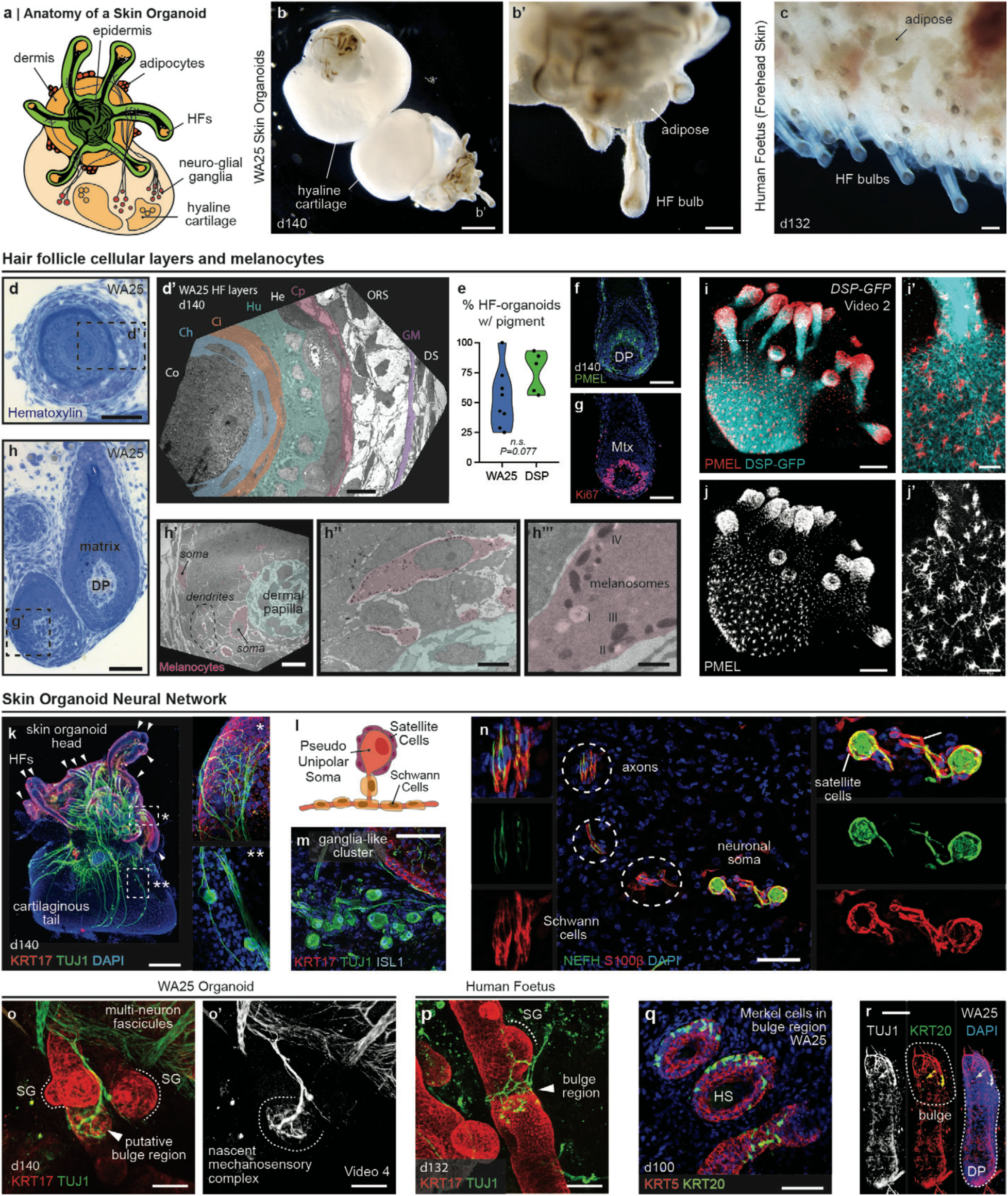
Induction of specialized cellular sub-types and a neural network in skin organoid cultures. **a**, Schematic of a typical skin organoid. The highlighted features are displayed in this figure and **Extended Data Figs. 7-9**. **b**, **c**, Comparison of pigmented HFs on (b, b’) day-140 WA25 skin organoids and (c) day-132 human foetal forehead skin. Note: adipocyte clusters that have formed in both samples. (b) Hyaline cartilage is visible in the tail region of the skin organoids. **d, d’, d)** Hematoxylin staining and (d’) TEM imaging showing that day-140 organoid HFs contain all of the cell layers known to exist in human follicles. Abbr: Dermal Sheath (DS); Glassy Membrane (GM); Outer Root Sheath (ORS); Companion (Cp) Layer; Henle’s (He) Layer; Huxley’s (Hu) Layer; Inner Root Sheath Cuticle (Ci,); Cuticle (Ch); Cortex (Co). Note: we were unsuccessful in capturing clear images of the medullar layer in our analysis. **e,** Violin plots comparing frequency of HF pigmentation in WA25 (53.5%, 70 pigmented hairy-organoids, n=137, 8 experiments) and *DSP-GFP* (76.2%, 72 pigmented hairy-organoids, n=91, 5 experiments) skin organoids (*p*=0.077). Abbr: not significant (n. s.). **f, g,** Immunostaining for (f) PMEL and (g) Ki67 reveals melanocytes integrating into a WA25 organoid HF matrix epithelium containing cycling cells. **h-h’’’,** (h) Hematoxylin staining and (h’-h’’’) TEM imaging showing matrix-associated melanocytes containing melanosomes. (h’’’) Melanosomes at different developmental stages (I, II, III, and IV) are visible. **i-j’,** Wholemount immunostaining for PMEL on *DSP-GFP* skin organoid shows concentrated melanocytes in the HF matrix and evenly distributed melanocytes in epidermis (see **Supplementary Video 2**). **k**, Wholemount immunostaining for KRT17 and TUJ1 reveals a day-140 WA25 skin organoid’s neural network. Insets (*, **) highlight neurite targeting to HF bulge region and neuron cell soma. Arrowheads indicates individual HFs. **l,** Schematic showing the typical pseudo-unipolar morphology of skin organoid neurons, which is consistent with large-to-medium sized cell soma cranial or dorsal root ganglion morphologies. **m,** Immunostaining for TUJ1 and ISL1 on day-140 WA25 organoid reveals the ganglia-like cluster of neurons. **n**, A subset of organoid neurons express Neurofilament-Heavy Chain and are associated with S100β-positive satellite glial cells and Schwann cells. **o-p,** Wholemount immunostainings for TUJ1 and KRT17 on (o, o’) day-140 WA25 skin organoid and (p) day-132 human foetus forehead tissue shows neurons innervating the HF bulge region (see **Supplementary Video 4**). Note: sebaceous glands are located superficial to the HF bulge region, consistent with normal hair structure. Arrowheads indicates HF bulge region. **q, r,** Wholemount immunostaining for KRT20 reveals Merkel cells in HF bulge region of day-100 WA25 skin organoid. (r) TUJ1^+^ neurons are targeting HF bulge region where KRT20^+^ Merkel cells are. All dashed boxes indicate the magnified area presented in the following images. Scale bars, 250 µm (**b, k**), 100 µm (**b’, c, i, j**), 50 µm (**f**, **g, h, m-r**), 25 µm (**d, i’, j’**), 10 µm (**d’, h’**), 5 µm (**h’’**), 800 nm (**h’’’**). Additional data in Extended Data Figs. 7 and 8.

Next, we investigated whether the neuro-glial progenitor cells identified by scRNA-seq at day 29 had matured into organized nervous tissue. In 100% of the organoids examined (>30 organoids from 7 experiments), we discovered a network of fasciculated βIII-Tubulin (TUJ1)^+^ Neurofilament-Heavy Chain (NEFH)^+^ axons interwoven between HFs (Fig. 3k, **Supplementary Video 2, 4**). The organoid neurons were reminiscent of neural crest-derived neurons found in cranial or dorsal root nerve ganglia that form during development. Their morphology was consistent with medium-large soma pseudo-unipolar neurons (Fig. 3k-n). We found that the neuron axons form fascicles that were associated with S100β^+^ Schwann-like cells. S100β^+^ satellite glia-like cells cover the neuron cell bodies, similar to cranial ganglia in human embryos (Fig. 3n). Neuronal processes appeared to contact the skin organoid epithelium and wrap around the circumference of the HFs, much like HFs in the day-132 human foetus (Fig. 3k, o, p, **Supplementary Video 5**). We determined that KRT20^+^ Merkel cells were located in the outer root sheath of HFs near the site of axon targeting, which we presume to be the bulge region of the HFs (Fig. 3q, r, **Supplementary Video 5**). Our data suggest that skin organoids may be able to form the diverse array of nerve endings found in normal skin—in particular, mechanosensitive touch complexes^35,36^. For confirmation, however, additional molecular and physiological profiling of the neural network will be necessary.

After ∼150 days in culture, we observed a build-up of dead squamous cells in the core of skin organoids (Extended Data Fig. 9b) and abnormal HF morphologies, suggesting that this timepoint may be the upper limit for maintaining skin organoids *in vitro*. Thus, we tested whether skin organoids could integrate into endogenous skin in a mouse model (Fig. 4a). On day 140, we removed the cartilaginous tails with tungsten needles and implanted hair-bearing WA25 skin organoids in small 1-2-mm incisions in the back skin of nude (NU/J) mice (Fig. 4b)^37^. Remarkably, after a month, we observed hairs growing out from 55% of the xenografts (15 of 27 xenografts, 3 experiments; Fig. 4c-e). The other grafts had in-grown hairs (22%) or failed (22%) (Fig. 4c). For out-grown xenografts, histology confirmed that the organoid epidermis had integrated with the host epidermis and the hair shafts were oriented perpendicular to the skin surface, reminiscent of bona fide human skin tissue (Fig. 4f, g). We noted that the xenograft epidermis had a cornified layer and rete ridge-like structures comparable to adult facial skin (Fig. 4f-h). In addition, CD49F^+^ vasculature had grown into the xenografts (Fig. 4g’). Our findings demonstrate that cystic skin organoids can unfurl and integrate into planar skin at a wound site.

**Figure 4.**
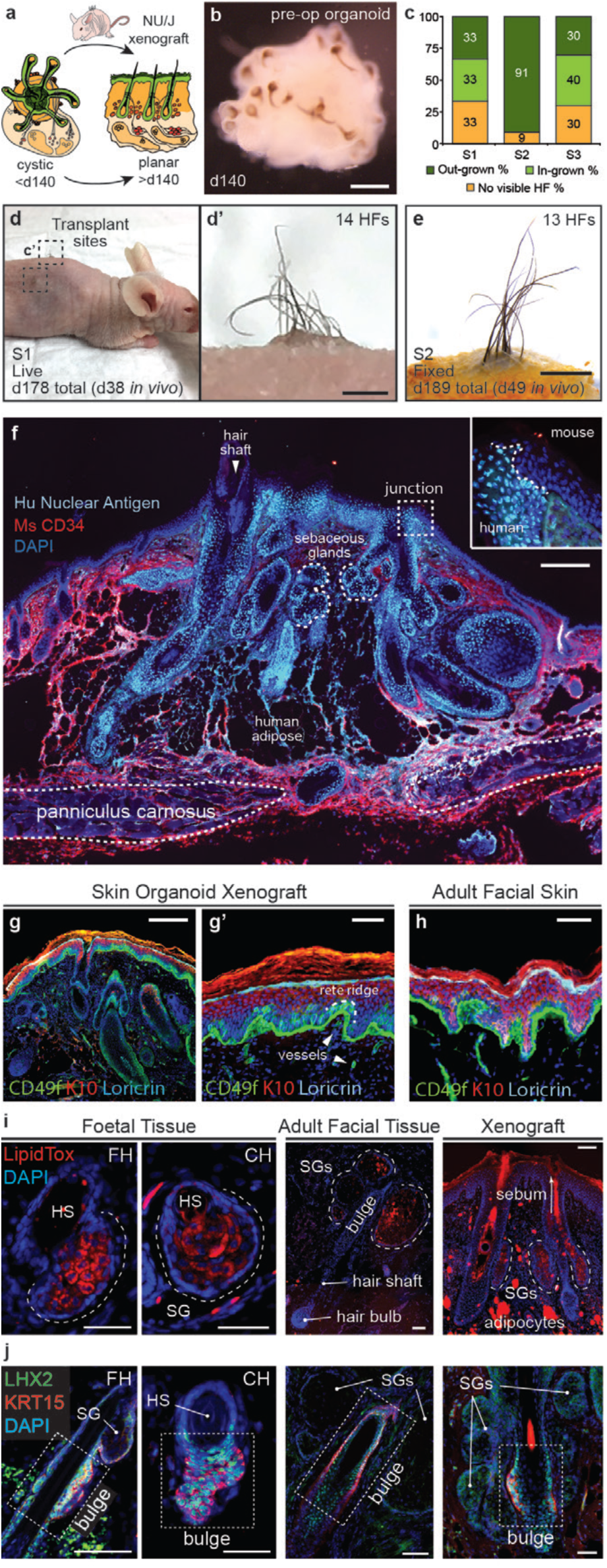
Skin organoids undergo cystic-to-planar transition to form hair-bearing skin in a nude mouse xenograft model. **a**, Schematic of skin organoid xenografting strategy. **b,** Representative day-140 WA25 skin organoids prior to grafting. **c**, Quantification of xenograft experiments. Data were compiled from 27 xenografts performed over 3 separate experiments/surgeries (S1, S2, S3). **d**, **d’**, A xenografted NU/J mouse 38 days post-operative (PO) following the first experiment (S1). Pigmented hair shafts are visible at both graft sites. **e**, An S2 xenograft containing out-grown pigmented hairs. **f**, Immunostaining of an out-grown xenograft showing the border between mouse and human tissue. Mouse connective tissue and dermis is labelled with a mouse-specific CD34 antibody. The human tissue has a been immunostained for human nuclear antigen. Note the seamless junction point between mouse and human epidermis. **g**-**h**, Comparison of an organoid xenograft (g, g’) to adult facial skin (h). **i**, Comparison of LipidTOX stained sebaceous glands in foetal tissue, adult facial skin, and an organoid xenograft. **j**, Comparison of putative bulge stem cells (LHX2^+^ KRT15^+^ cells) in foetal tissue, adult facial skin, and an organoid xenograft. Scale bars, 1 mm (**d’, e**), 200 µm (**a, f, g**), 100 µm (**i**, **j; Adult Facial tissues, Xenografts**), 50 μm (**g’**, **h**, **i, j; Foetal Tissues**). Additional data in Extended Data Fig. 9 and 10.

Finally, we examined out-grown xenografted HFs for two hallmarks of mature pilosebaceous units: sebaceous glands and bulge stem cells. We detected sebaceous glands containing SCD1^+^ cells in all xenografted HFs (Fig. 4i, Extended Data Fig. 9e-f). Organoid sebocytes had sebum vacuoles that were visible under electron microscopy (Extended Data Fig. 9d). Morphologically, xenograft sebaceous glands appeared to have multiple lobes similar to adult sebaceous glands (Fig. 4i, Extended Data Fig. 9). The sebaceous glands are typically located superficial to the bulge region of the HF. In the presumptive bulge region of xenografted HFs, we found KRT15^+^ NFATC1^+^ HF stem cell-like cells and Nephronectin (NPNT)^+^ basement membrane (Fig. 4j, Extended Data Fig. 10d**, Supplementary Video 6**). Notably, the expression pattern of NFATC1^+^ was predominantly cytoplasmic, which was consistent with follicular bulges in foetal skin (Fig. 4j). By contrast, adult bulge cells displayed nuclear NFATC1 expression (Extended Data Fig. 10)^38^. Together these findings suggest that skin organoid follicles reach a level of maturity equal to or beyond that of second-trimester facial hair. Long-term (>1 year) observation will be needed to determine if xenografted follicles can initiate and maintain a growth cycle.

In summary, our study establishes a method for generating skin organoids from PSCs; thus, providing a novel model with which to investigate the cellular dynamics of developing human skin^3,6^. Numerous genetic skin disorders and cancers could be modelled with skin organoids to accelerate drug discovery^39^. Moreover, with additional control of cell lineage specification, we envision that skin organoids could be used to reconstitute appendage-bearing skin in burned or wounded patients.

## ACKNOWLEDGEMENTS

This work was supported by the Ralph W. and Grace M. Showalter Trust (K.R.K.), the Indiana CTSI (core pilot grant UL1 TR001108 to K.R.K.), the Indiana Center for Biomedical Innovation (Technology Enhancement Grant to K.R.K.), and the NIH (grant DC015624 to K.R.K.). Cell lines associated with this study were stored in a facility constructed with support from the NIH (grant C06 RR020128-01). The University of Washington Birth Defects Research Laboratory was supported by NIH award number 5R24HD000836 from the Eunice Kennedy Shriver National Institute of Child Health and Human Development. We would like to thank B. Koh, A. Tward, S. Frumm, D. Spandau, J. Foley, U. Arimpur, E. Longworth-Mills, P-C. Tang, A. Elghouche, M. Kamocka, and C. Miller for their technical assistance. We especially thank M. Rendl and N. Saxena for critical comments on the manuscript.

## AUTHOR CONTRIBUTIONS

J.L. and K.R.K. conceived the study and wrote the manuscript. J.L. performed most of the *in vitro* experiments and IHC analysis. J.L., C.R., Z.P., T.S., and K.R.K. designed and performed the *in vivo* experiments. M.S. and A.K. performed IHC and scRNA-seq data analysis. H.G. and Y.L. performed bioinformatic analyses of scRNA-seq data and generated figures. B.M.W. and S.H. performed protocol validation experiments and generated figures. K.R.K. supervised the project, monitored the experiments, and acquired funding. All authors read and approved the final manuscript.

## METHODS

No statistical methods were used to predetermine sample size. The experiments were not randomized, and the investigators were not blinded during experiments and outcome assessments.

### hPSC lines and culture

Culture experiments were performed with the WA25 hESC line (passage 21-47), purchased from WiCell Research Institute, and the Desmoplakin-mEGFP (*DSP-GFP*) hiPSC line (passage 35-51), acquired from the Allen Institute for Cell Science and the Coriell Institute^40^. Cells were cultured on 6-well plates coated with Vitronectin Recombinant Human Protein (Invitrogen) at a concentration of 0.5 µg/cm^2^. Pluripotent stem cells were maintained in Essential 8 Flex (Gibco) medium with 100 µg/ml Normocin (Invivogen; hereafter, E8). The medium was replenished every other day. Cells were passaged at ∼80% confluency (generally every 4-5 days) in medium containing 10 µM Y27632 (hereafter, Y; ROCK inhibitor that inhibits apoptosis, ReproCell). WA25 cells were passaged in clusters with 0.5 mM EDTA, and *DSP-GFP* cells were passaged as single cells with StemPro Accutase Cell Dissociation Reagent (Gibco). Fresh E8 medium was replenished twenty-four hours following passage. Detailed information concerning cell line specifics as well as validation and testing is available at: https://www.wicell.org/home/stem-cells/catalog-of-stem-cell-lines/wa25.cmsx?closable=true and https://www.allencell.org/cell-catalog.html.

### Optimized hPSC differentiation

For differentiation, pluripotent hPSC colonies were detached from the culture dish using StemPro Accutase (Invitrogen). Once dissociated, hPSCs were collected as a single cell suspension in E8 medium containing 10 µM Y (Stemgent; hereafter, E8-10Y). Cell concentration was determined with an automated counter (Countess II, Invitrogen) with Trypan Blue. The appropriate number of cells needed was transferred into E8 medium containing 20 µM Y (hereafter, E8-20Y). From the E8-20Y cell suspension, cells were distributed at a count of 3,500 cells in 100 µl per well into 96-well U-bottom plates. Aggregation was aided by centrifugation at 110 rcf for 6 min. These cell aggregates were incubated in 37°C incubator under 5% CO_2_ for 48 hrs. Thus, we designated this timepoint as day −2. After 24 hrs incubation (at day −1), 100 µl E8 medium was added to each well in order to dilute out Y and promote cell proliferation and aggregate growth. On day 0, to **start differentiation**, all cell aggregates were collected and transferred to a new 96-well U-bottom plates in 100 µl of E6-based differentiation medium containing 2% Matrigel (Corning), 10 µM SB (Stemgent), 4 ng/ml basic-FGF (hereafter, FGF2; PeproTech), and 2.5 ng/ml BMP-4 (PeproTech) to initiate non-neural ectoderm formation. On day 3 of differentiation, to induce CNCC formation, 200 ng/ml LDN, a BMP inhibitor, and 50 µg/ml of FGF2 were added in a volume of 25 µl per well, thus making the final volume 125 µl per well. On day 6 of differentiation, 75 µl of fresh E6 medium was added, bringing the final volume to 200 µl. Half of the media was changed (removal of 100 µl spent medium and addition of 100 µl fresh E6 medium) on days 8 and 10. On day 12, in order to induce self-assembly of epithelium, all aggregates were transferred into individual wells of a 24-well low-attachment plate in 500 µl organoid maturation medium (OMM) containing 1% Matrigel. To maintain the aggregates in a floating culture for constant medium circulation, 24-well plates were placed on an orbital shaker at 65 rpm in the 37°C incubator with 5% CO_2_. OMM is Advanced DMEM/F12 (Gibco) and Neurobasal (Gibco) media at a 1:1 ratio, 1X GlutaMax™ (Gibco), 0.5X B-27 Minus Vitamin A (Gibco), and 0.5X N2 (Gibco) supplements, 0.1 mM 2-Mercaptoethanol (Gibco), and 100 µg/ml Normocin (Invivogen). On differentiation day 15, half of the spent medium was replenished (removal of 250 µl of spent medium and addition of 250 µl fresh medium) with OMM containing 1% Matrigel. Starting from day 18, half-medium was changed every three days (from day 18 to day 30) or every other day (from day 30 to day 150 or longer) with fresh OMM without Matrigel. Increasing total volume of medium per well to 1 ml was at times necessary from day 80 onward as aggregates mature and grow larger. (See **Supplementary Note** for additional differentiation protocol information and **Supplementary Table 2** for media compositions).

### Human Foetal and Adult Specimens

Facial tissue samples from miscarried foetuses were obtained from the University of Washington Birth Defects Research Laboratory. The tissue was not obtained from living individuals, and was de-identified. This work falls under NIH Exemption 4, thus it carries a “Non-Human Subjects Research” status by the Indiana University School of Medicine IRB. De-identified adult human facial skin tissue samples were obtained from the Human Skin Disease Resource Centre at Brigham and Women’s Hospital-Harvard Medical School.

### Skin organoid dissociation for scRNA-seq

Five representative skin organoids for each condition (day-29 WA25 and *DSP-GFP*) were pooled for cell dissociation. For day-48 WA25 cell aggregates, the tail structure containing non-skin associated mesenchymal and neuronal cells was removed using two tungsten needles prior to dissociation in order to bias analysis toward skin tissue. Briefly, collected organoids were incubated with pre-warmed (37°C) TrypLE™ for 30 min in 37°C incubator on a shaker at 65 rpm for a gentle swirl. During 30 min of incubation, the dissociation mixture was agitated every 10 min with gentle pipetting using wide-bore p1000 and p200 tips (Mettler-Toledo). By 30 min of incubation, the tissue structure dissociated into a single cell suspension with no visible cell aggregation. Cold 3% BSA solution (Sigma-Aldrich) was added to the dissociated cell suspension in order to inactivate TrypLE enzymatic activity. The suspension was filtered through a 40 µm Flowmi™ cell strainer (Bel-Art™) in order to eliminate any debris. An additional three washes with cold 3% BSA solution were performed. Cells were resuspended in 3% BSA, and filtered through a 40 µm Flowmi™ cell strainer. Viability (live cell percentage) and live cell count were determined using Trypan Blue (Gibco; used at 1:1 ratio) and Countess™ II Automated Cell Counter (Life Technologies). Final cell concentration was ∼1,000 cells/µl and >90% viability.

### scRNA-seq cDNA library preparation and sequencing

Single cell 3’ RNA-seq experiments were conducted using the Chromium single cell system (10x Genomics, Inc) and the NexSeq 500 sequencer (Illumina, Inc). Approximately 10,000 cells per sample were added to a single cell master mix, following the Chromium Single Cell 3’ Reagent Kits v2 User Guide, CG00052 Ver B (10x Genomics, Inc). Along with the single-cell gel beads and partitioning oil in separate wells of a Single Cell A Chip, the single cell reaction mixture was loaded to the Chromium Controller for GEM generation and barcoding, followed by cDNA synthesis and library preparation. At each step, the quality of cDNA and library was examined by Bioanalyzer. The resulting library was sequenced in a custom program for 26b plus 98b paired-end sequencing on an Illumina NextSeq 500 to a read depth of >30,000 reads per cell.

### scRNA-seq data analysis

The 10x Genomics CellRanger 2.1.0 pipeline (http://support.10xgenomics.com/) was used to process raw sequence data. Briefly, Cellranger uses bcl2fastq (https://support.illumina.com/) to demultiplex raw base sequence calls generated from the sequencer into sample-specific FASTQ files. The FASTQ files are then aligned to the reference genome with RNAseq aligner STAR. The aligned reads are traced back to the individual cells and the gene expression level of individual genes are quantified based on the number of UMIs (unique molecular indices) detected in each cell. Filtered gene-cell barcode matrices were generated by CellRanger for further analysis. Cells with extremely high or low number of detected UMIs were excluded from further analysis. In addition, cells with a high percentage of mitochondrial reads were filtered out. After removing unwanted cells, the gene expression levels for each cell were normalized by the total number of UMIs in the cell and multiplies by a scaling factor of 10,000. After log-transformation, we used Seurat 2 for cell clustering using principle component analysis (PCA) on highly variable genes^41^. The cell clusters were visualized using the T-Distributed Stochastic Neighbor Embedding (t-SNE) and Uniform Manifold Approximation and Projection (UMAP) plotting methods. The gene markers for each cluster were identified through differential expression analysis by comparing cells in the cluster to all other cells (x.low.cutoff=0.0125, x.high.cutoff=4, y.cutoff=0.5). For most gene-specific tSNE or UMAP plots throughout the manuscript were generated using x.low.cutoff=0.25 and x.high.cutoff=2.

To integrate the single cell data from WA25 and DSP-GFP samples, we applied the canonical correlation analysis (CCA) in Seurat^42^. We chose the top 1500 variable genes from each sample to calculate the correlation components (CCs) and used the function MetageneBicorPlot to determine the optimal number of CCs. We retained the cells whose expression profile could be explained with at least 50% by the CCs using CalcVarExpRatio and SubsetData. The CCA subspaces were then aligned with AlignSubspace using the number of CCs determined. We employed FindClusters for shared nearest neighbor (SNN) graph-based clustering. The clusters were visualized with t-distributed stochastic neighbor embedding (t-SNE) by running dimensionality reduction with RunTSNE and TSNEPlot. The FindConservedMarkers function was subsequently used to identify canonical cell type marker genes that are conserved of cells across different samples. To compare average gene expression within the same cluster between cells of different samples, we applied function AverageExpression. R packages ggplot2 and ggrepel (https://github.com/slowkow/ggrepel) were used to plot the average gene expression. Violin plots (VlnPlot) and feature plots (FeaturePlot) were used to visualize specific gene expressions across clusters and different sample conditions.

In the supplementary material we have provided four HTML files that generate an online interface with which to explore our scRNA-seq analysis pipeline and evaluate additional cell cluster markers (**Supplementary Data 1-4 *will be available with the peer-reviewed version of the manuscript***). Additionally, the raw sequencing files and 10x CellRanger output files will be provided on the Gene Expression Omnibus. For quick visualization and exploration of cell clusters in our datasets we recommend using the 10x Genomics Loupe Browser or the SPRING online portal^43^ (kindly hosted by Allon Klein, Harvard Medical School): https://kleintools.hms.harvard.edu/tools/spring.html.

### Immunohistochemistry

For immunostaining, fixed samples were cryoprotected through a graded treatment process of 10, 20, and 30% of Sucrose (Sigma-Aldrich), embedded on cryomolds (Endwin Scientific) in tissue freezing medium (General Data Healthcare), snap frozen at −80°C, and sliced to 12 or 15 µl thickness. Cryosections were blocked in 10% normal goat/horse serum, incubated with primary antibodies diluted in 3% normal goat/horse serum, and then incubated with secondary antibodies in 3% normal goat/horse serum. Unless stated otherwise, images are representative of specimens obtained from at least 3 separate experiments. For IHC analysis of aggregates between days 0-12, we sectioned 3-6 aggregates from each condition in each experiment. IHC analysis of later stages of development was performed on at least 3 aggregates from each condition per experiment.

Whole-mount staining was performed with a previously published method^43^ with major adjustments in incubation times to account for reduced tissue size^4^. In brief, samples were fixed in 4% (v/v) PFA (Electron Microscopy Sciences), permeabilized, descaled, incubated with primary antibodies, incubated with fluorescently-labelled secondary antibodies, re-fixed in 4% (v/v) PFA (Electron Microscopy Sciences), and cleared. Microscopy was performed with a Leica DMi8 Inverted Microscope, a Leica TCS SP8 Confocal Microscope, or an Olympus FV1000-MPE Confocal/Multiphoton Microscope. Three-dimensional reconstruction was performed with Imaris 8 software package (Bitplane) on computers housed at the Indiana Centre for Biological Microscopy. See **Supplementary Table 3** for a list of antibodies.

### Cell culture quantitative analysis: Frequency of organoids producing HFs

Starting from day 75, organoids with protruding hair placodes, germs, pegs, and HFs were counted and recorded. To calculate a percentage of HF production within a culture, number of HF-produced organoids was divided by total number of organoids cultured in the experiment and multiplied by a hundred. Percentages of HF formation from nine independent experiments of both WA25 and *DSP-GFP* cell lines were calculated and averaged to get the general frequency of HF production from each cell line.

#### Number of HFs produced

HF numbers generated in both WA25 and *DSP-GFP* skin organoids were counted manually. From each cell line-derived hairy skin organoid in nine independent experiments, 80 organoids were randomly pooled, and pictures were taken under Leica M165FC stereo microscope. An angle of each organoid was chosen blinded, and a picture of that view under the microscope was taken. Number of HFs in the image view was quantified by counting the protruded hair bulbs using ImageJ ‘Cell Counter’ plugin function. Interestingly, the bigger skin organoid cyst is, the higher number of HFs is produced.

#### Frequency of HF pigmentation

Among the hair-bearing skin organoids, some HFs were pigmented in dark brown or black in the bulbs and hair shafts, while the rest were albino – white hairs. Pigmentation is visible with naked eyes, but to clearly distinguish pigmented hairs, including faint, pigmentation-initiating HFs from white hairs, organoids were observed under Leica DMi1 without condenser on and/or under Leica M165FC stereo microscope set to dark field illumination. Number of skin organoids containing pigmented HFs was divided by the total number of HF generating organoids within the experiment and multiplied by a hundred to calculate the percentage of pigmented HF formation. Each percentage from eight independent WA25 experiments and five independent *DSP-GFP* experiments were averaged to determine the HF pigmentation rate of each cell line.

### LipidTOX Staining

LipidTOX Staining (Invitrogen) was performed to visualize sebaceous gland and lipid-rich adipocytes. Cryosections of skin organoids were incubated with LipidTOX neutral lipid stain diluted at a ratio of 1:200 in 1X PBS for 30 min at RT, followed by Hoechst staining (1:2000; Invitrogen) for 1 min at RT to visualize nuclei.

### Xenograft Experiments

*In vivo* experiments were performed after an approval from the Indiana University Animal Care and Use Committee (IACUC) and complied federal regulations. For grafting, 5-6 week-old female immunodeficient nude mice (NU/J) purchased from The Jackson Laboratory were used. Prior to the graft surgery, WA25 cell-derived either pigmented or albino hair-bearing skin organoids at varying days – days 113, 115, 120, 124, 127, 140, and 145 – were pooled and prepared; using tungsten needles, tail portion (mesenchymal cells or cartilages) of the organoids was gently removed, and a tiny incision was made on the hairy-skin to increase chances for the organoid core to be exposed and allow hairs to grow outward. The mice were anesthetized under Isoflurane, and Bupiviczaine, a pain reliever, was administered (6 mg/kg) via subcutaneous injection to the intended incision sites on the back of the mice (3-5 incision sites per mouse). About 1-2 mm-sized incisions, of which can fit a hairy-skin organoid, were made, and each hairy-skin organoid prepared ahead was placed on individual incision site by facing hair bulb side inward, contacting muscle layer of the mice, and organoid-incision side outward, being exposed to the air. Sterile gauze was place on top of the mice’s back, covering all graft sites, and tightened with bandage by wrapping around the body of the mice. Then, the bandage was sutured for stability. After the surgery, the mice were provided with Carprofen wet food for pain relief. Dressing was removed 7-9 days post-surgery. The mice were observed for- and sacrificed after-about 7 or 14 weeks of the surgery. Using a scalpel, each area with skin organoid graft was collected at about 3 mm x 5 mm (width x length) size. Collected samples were flattened on a filter paper to inhibit curling effect of the skin edges and fixed in 4% PFA overnight at 4°C with constant gentle agitation.

### Analysis of HFs growth in xenografts

For transplantation, skin organoids containing hairs were selected and grafted at approximately day 140 of differentiation. HF protrusion and growth at the grafted sites on the back of mice were monitored until the day mice were sacrificed. Some HFs failed to grow outwards potentially due to imperfect organoid grafting method that the follicle orientation was altered (hair bulbs of organoid facing outward) during the procedure that the hairs grew inward instead and were visible as pigmented dark shade under the skin. Number of grafted sites with protruded hairs (both pigmented and albino) or dark pigmented shade under the skin were counted, divided by the number of initial grafts that were made to calculate the percentage of out-grown, in-grown, or failed grafts.

### Statistical analysis

All statistics were performed using GraphPad Prism 8 software. Statistical significance was determined using an unpaired Student’s t-test with Welch’s correction to account for unequal standard deviations and sample sizes. Unless stated otherwise data are reported as mean ± SEM. No statistical test was used to predetermine sample size, the investigators were not blinded to the treatment groups, and the samples were not randomized.

### Protocol development and optimization (Koehler Lab)

The culture method was optimized and replicated by two investigators (K.R.K., and J.L.) using the WA25 and *DSP-GFP* cell lines.

### Protocol validation (Heller Lab)

During preparation of this manuscript, K.R.K. and J.L. contacted S.H. and B.M.W. for assistance validating the skin organoid culture method. S.H. and B.M.W. were provided with a detailed protocol for skin organoid induction with a list of suggested reagents. S.H. and B.M.W. were blinded to the data already generated for this study with WA25 and *DSP-GFP* PSCs; thus, they were unfamiliar with the expected morphological changes during skin organogenesis. K.R.K. and J.L. provided minimal guidance via email. In his initial experiment, B.M.W. generated 92 hair-bearing skin organoids (out of 130 total organoids) from WA01 (H1) ESCs (see Fig. 2g, h). An additional validation experiment is underway.

### Data availability

Data are available from the corresponding author upon reasonable request.

## FIGURES AND LEGENDS

**Extended Data Figure 1.**
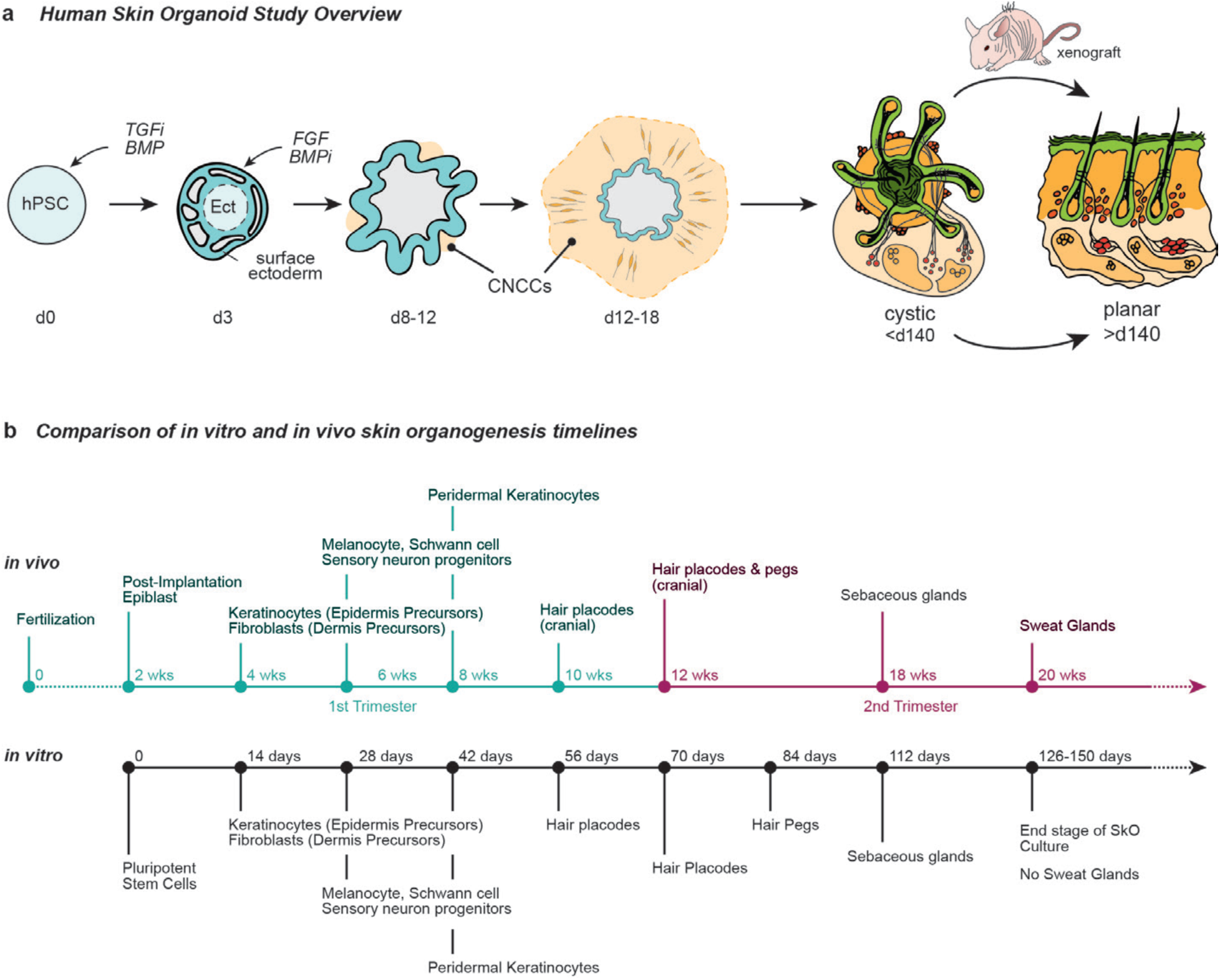
Overview of skin organoid induction in relation to normal human foetal skin developmental events. **a**, Overview of the skin organoid protocol. TGFi/BMPi = signalling pathway inhibition. Cranial Neural Crest Cell (CNCC). **b**, Comparison of *in vitro* and *in vivo* skin development timelines. Note that developmental timing is approximate. Corresponds with data/concepts in Fig. 1.

**Extended Data Figure 2.**
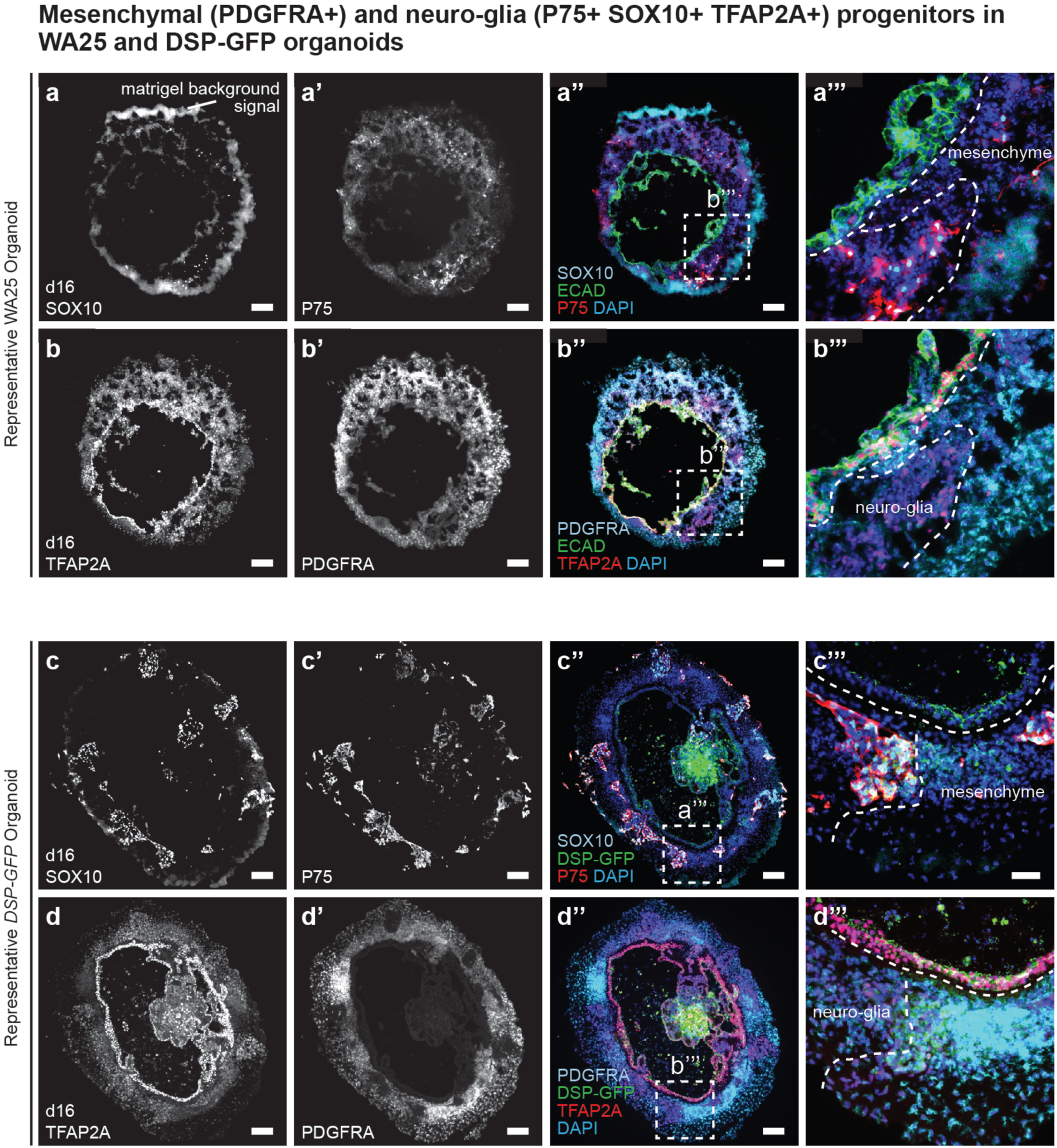
Surface ectoderm and CNCC induction in WA25 and *DSP-GFP* skin organoid cultures. **a, b**, Cryosectioned WA25 cell aggregates at day 16 of differentiation immunostained for SOX10, P75, TFAP2A, ECAD, and PDGFRα. ECAD (green) labels the epithelium, while SOX10/P75 and PDGFRα label distinct populations of presumptive neuro-glial and mesenchymal progenitors, respectively. **c, d,** Cryosections of *DSP-GFP* cell aggregates at day 16 of differentiation immunostained for the same markers as in panels (a, b). ECAD was omitted to allow observation of the endogenous DSP-GFP signal at the apical surface (see panels c’’’ and d’’’) of the epithelium. Scale bars, 100 µm (**a-a’’, b-b’’, c-c’’, d**-**d’’**), 50 μm (**a’’’, b’’’, c’’’, d’’’**). Corresponds with data in the top half of Fig. 1.

**Extended Data Figure 3.**
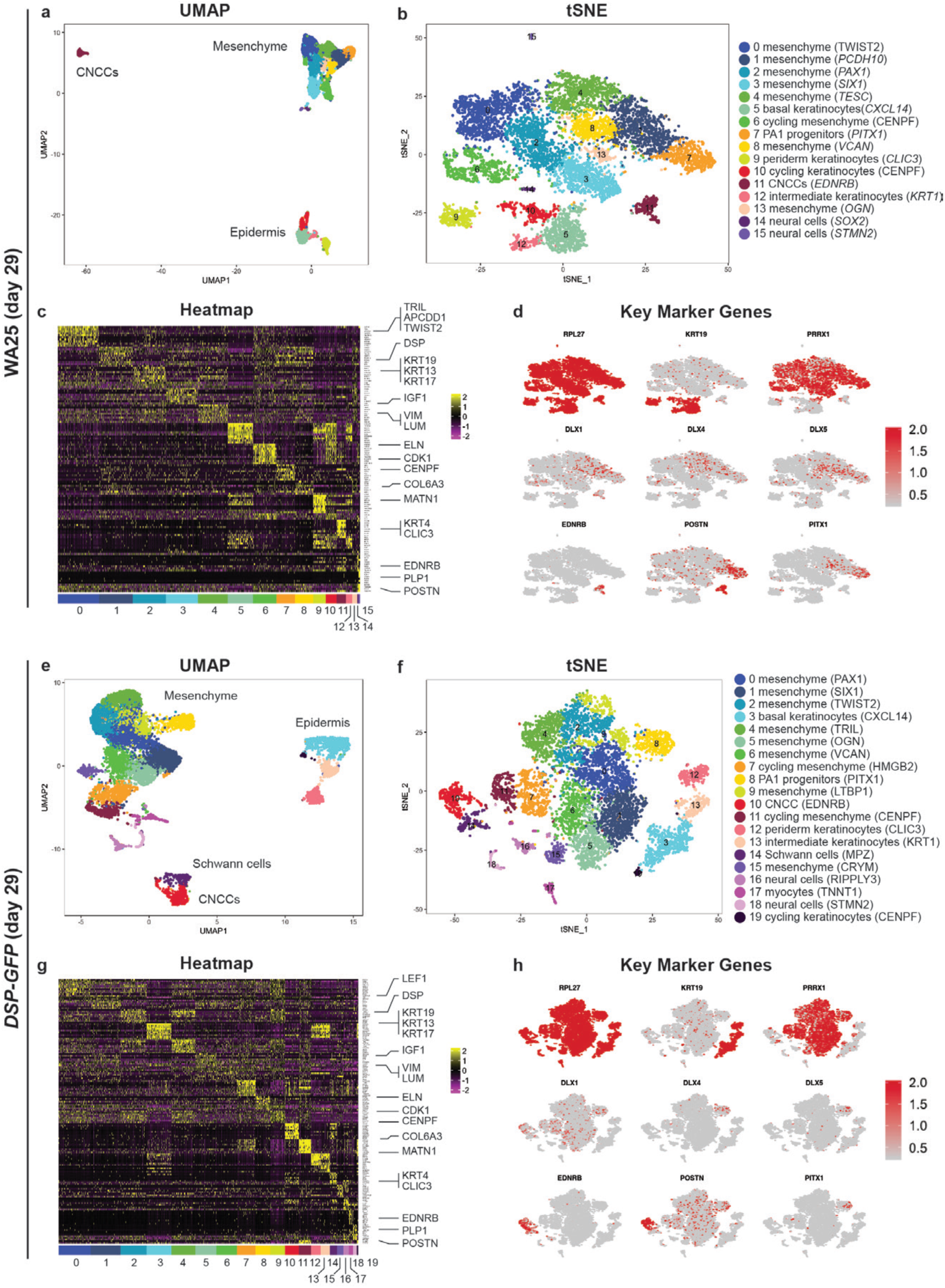
Single-cell RNA-seq analysis of day-29 skin organoids derived from WA25 and *DSP-GFP* cells. **a, e**, Uniform manifold approximation projections (UMAP) of WA25 and *DSP-GFP* cell clusters. The major cell cluster groupings of mesenchyme, epidermis, and CNCCs are noted. **b**, **f**, tSNE plots of WA25 and *DSP-GFP* cell clusters. **c**, **g**, Heatmap displaying the top 10 positively expressed genes per cell cluster for the day-29 WA25 and *DSP-GFP* scRNA-Seq datasets. **d, h,** UMAP plots for specific marker genes that define cell subtypes (*KRT19*, *PRRX1*, *EDNRB*, *POSTN*) and indicate pharyngeal arch identity (*DLX1*, *DLX4*, *DLX5, PITX1*). *RPL27*, a ribosomal housekeeping gene, is included as a positive control. Corresponds with data in the bottom half of Fig. 1.

**Extended Data Figure 4.**
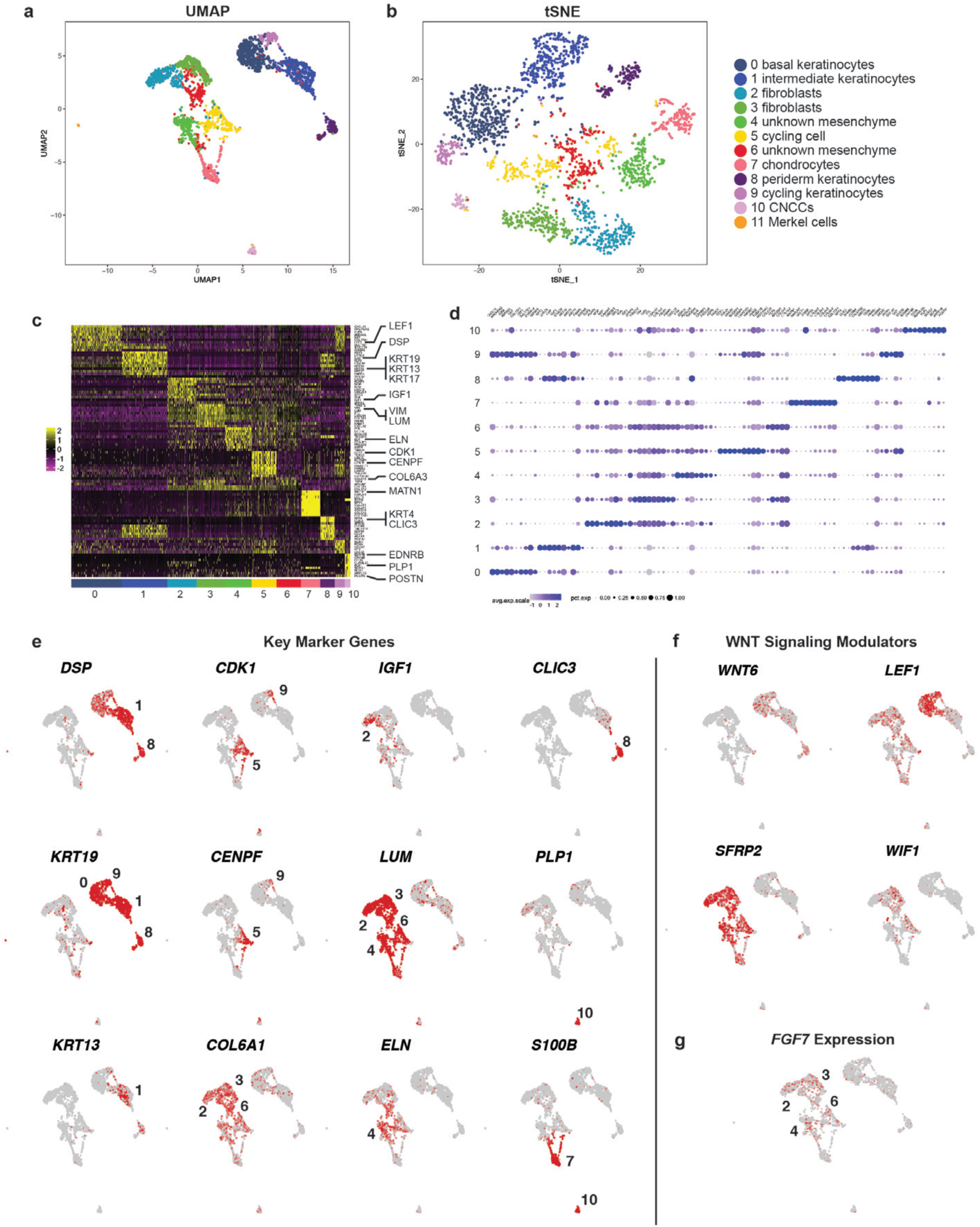
Single cell-RNA-seq analysis of day-48 skin organoids derived from WA25 cells. **a**, UMAP of day-48 WA25 cell subtype clusters. Using the UMAP clustering algorithm, we identified a subset of 9 cluster (C)-0 cells (putative basal keratinocytes) that were completely separated from the majority of C-0 cells, suggesting that our unbiased analysis pipeline failed to identify a unique subset of low-abundance cells. We used the Seurat manual selection tool to generate an 11^th^ cluster containing these cells. Among the statistically significant positive expressed genes in C-11 cells we identified *ATOH1*, *ISL1*, *SOX2*, *KRT8*, *KRT18*, and *KRT20,* which strongly suggest a Merkel cell identity. **b,** tSNE plot of day-48 WA25 cell clusters. **c**, Heatmap displaying the top 10 differentially expressed genes per cell cluster for the day-48 WA25 scRNA-Seq dataset. **d**, Dot plot array displaying the top 10 positively expressed genes per cell cluster for the day-48 WA25 scRNA-Seq dataset. **e,** UMAP plots for cell subtype specific marker genes. The numbers of cell cluster with positive expression are listed on the UMAP plot. **f**, UMAP plots for WNT signalling pathway genes. Note that *WNT6* is expressed in basal keratinocytes and peridermal keratinocytes. *LEF1* expression appears localized to basal keratinocytes. Negative WNT modulatory genes, *SFRP2* and *WIF1*, are expressed in the mesenchymal cell group. Corresponds with data in Fig. 2.

**Extended Data Figure 5.**
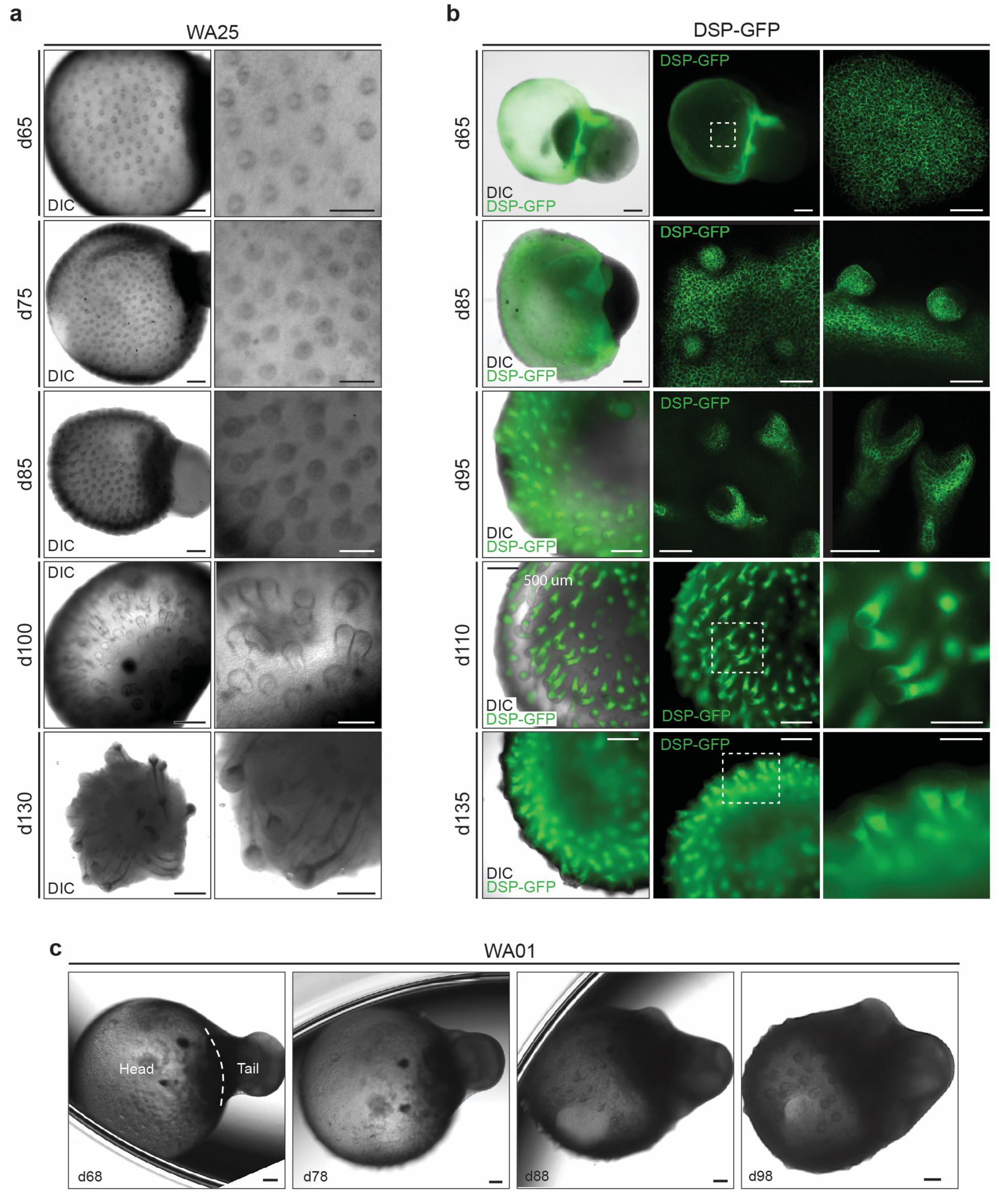
Comparison of initial follicle induction in WA25 and *DSP-GFP* live-cell aggregates. **a**, Differential interference contrast (DIC) images of d65-130 WA25 skin organoids with developing HFs. **b**, DIC and endogenous GFP fluorescence images of d65-135 *DSP-GFP* skin organoids with developing HFs. **c**, DIC images of d68-98 WA01 skin organoids with developing HFs. Scale bars, 500 µm (**a**-left column, **b**- left/middle column), 250 µm (**a**-right column, **b**-right column, **c**, all panels), 100 µm (**b**- d65/85 right column). Corresponds with data in Fig. 2.

**Extended Data Figure 6.**
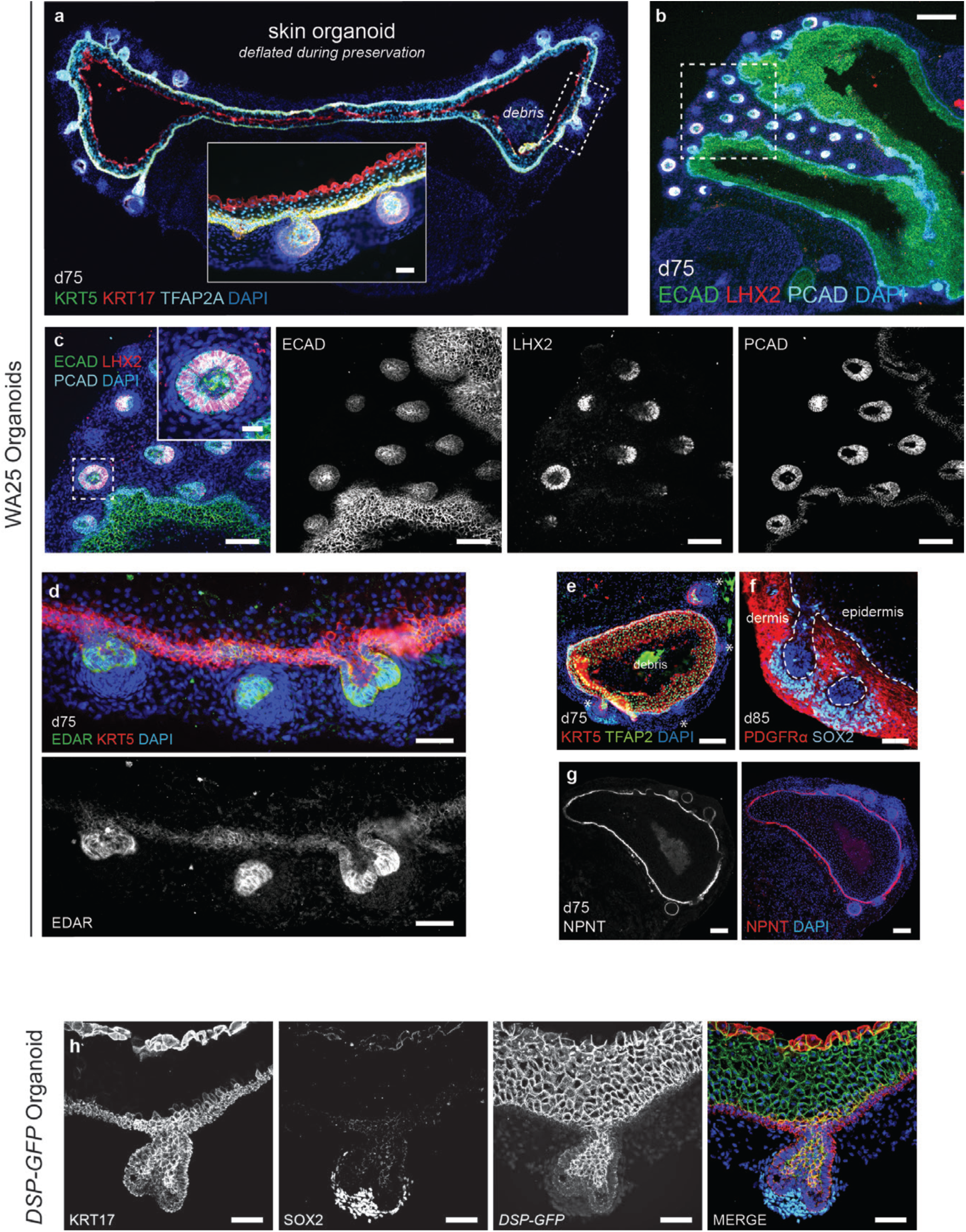
Key protein markers of HF induction in skin organoids. **a**, Day-75 WA25 skin organoid with nascent HFs immunostained for KRT5, KRT17, and TFAP2A. Note that this KRT17 antibody labels basal and peridermal keratinocytes. Dashed box region is presented in a high-magnification image. **b**, ECAD labels the entire epithelium, whereas PCAD expression is restricted to the basal layer and the hair germ epithelium. LHX2 labels hair placode and matrix cells. Dashed box indicates magnified region presented in panel (c). **c**, Higher magnification images of panel (b). Close up hair germ image in the dashed box region is presented on the right top corner of the merged image. Immunostainings of each marker are presented in separate channels. **d,** EDAR is also expressed in HF placodes and matrix cells. **e**, Low magnification image of a day-75 WA25 organoid showing the distribution of KRT5 and TFAP2A expression. **f**, PDGFRα is expressed throughout the outer layer of dermal cells. SOX2 labels dermal condensate and derma papilla cells, yet is also expressed in some basal epidermal cells—likely Merkel cells. **g**, NPNT (Nephronectin) is localized to the basement membrane of day-75 skin organoid epithelia. **h**, Separate channels for the *DSP-GFP*, KRT17, SOX2 immunostaining image shown in Figure 2n-n’’. Scale bars, 250 µm (**b**), 100 µm (**c, e, g**), 50 μm (**a-inset**, **d**, **f, h**). Corresponds with data in Fig. 2.

**Extended Data Figure 7.**
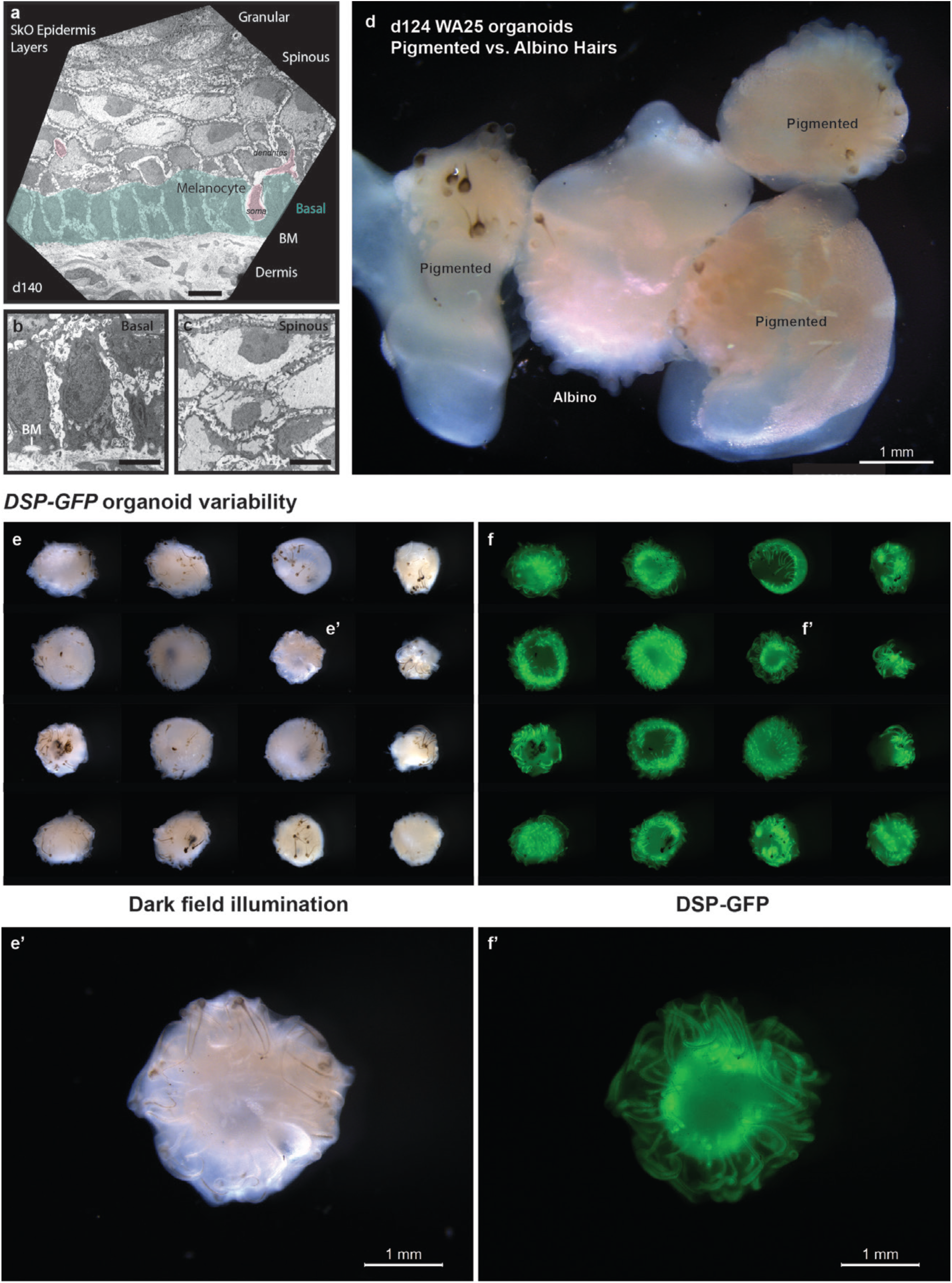
Pigmentation in WA25 and *DSP-GFP* skin organoid HFs. **a**, TEM image of epidermis in a day-140 WA25 skin organoid. A melanocyte cell body is pseudo-coloured pink, and basal skin layer is pseudo-coloured green. **b**, Higher magnification image of basal keratinocytes and the basement membrane (BM). **c**, Higher magnification image of spinous layer keratinocytes. **d,** Dark field illumination image of day-124 WA25 skin organoids, comparing pigmented vs. non-pigmented (albino) hairs development. **e-f’**, Overview of one *DSP-GFP* experiment containing 24 skin organoids. Note that each organoid displays pigmented HFs; however, the morphology of the epidermal cyst, as shown by DSP-GFP expression (green), varies between organoids. Scale bars, 1 mm (**d**, **e’**, **f’**), 10 µm (**a**), 5 µm (**b, c**). Corresponds with data in Fig. 3.

**Extended Data Figure 8.**
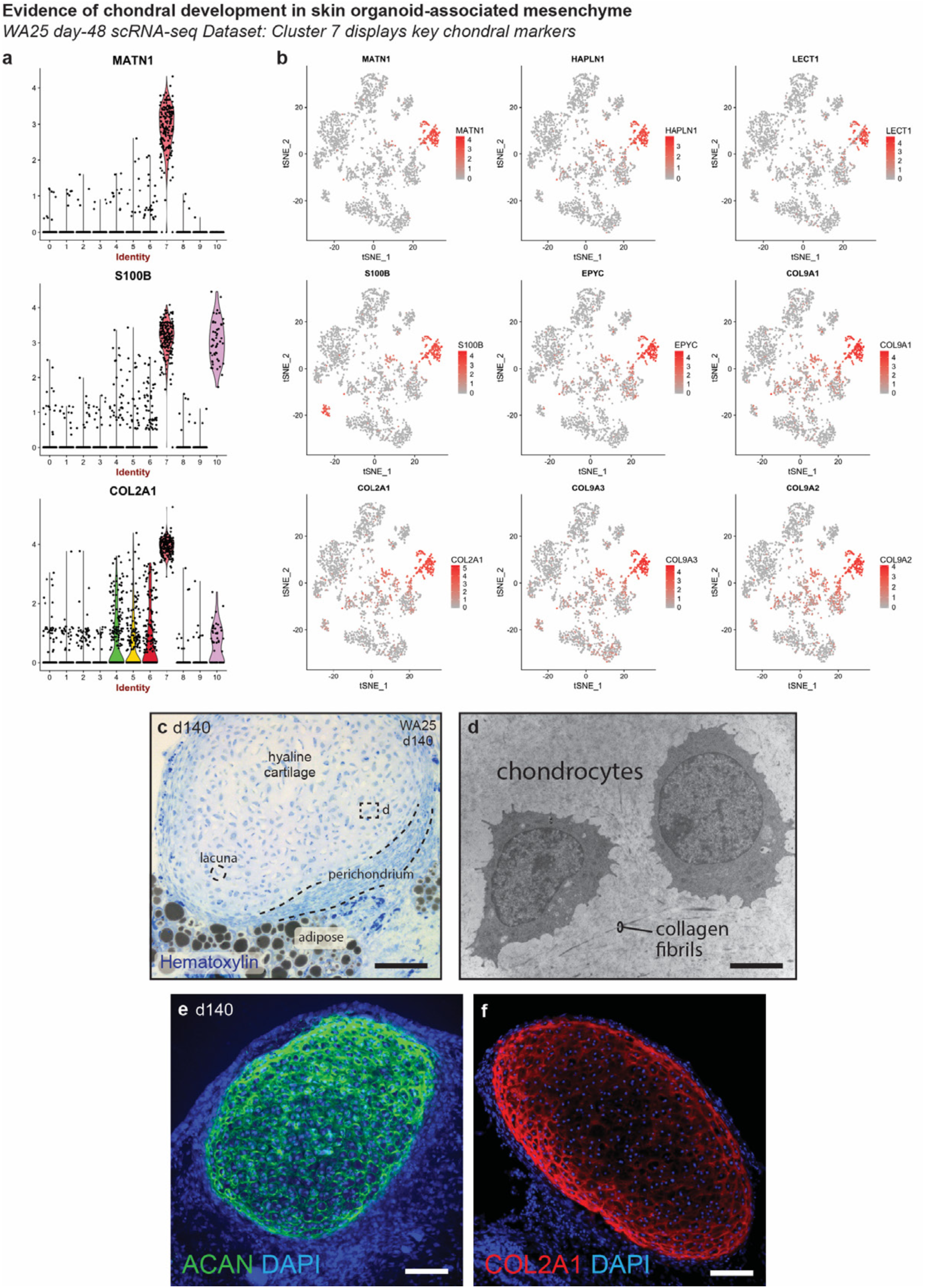
Chondral development in the skin organoid tail region. **a, b**, Violin and tSNE plots showing expression of chondral marker genes within cluster 7 of the day-48 WA25 skin organoid dataset (see Fig. 2 and **Supplementary Data 4**). **c**, Hematoxylin stained section showing hyaline cartilage that has formed within skin organoid-associated mesenchymal tissue (day 140). **d**, TEM image of two representative chondrocytes located within hyaline cartilage tissue shown in panel (c). **e, f,** Immunostaining of day-140 organoid samples for Aggrecan (ACAN) and Collagen 2A1 (COL2A1) highlights cartilage development. Scale bars, 100 μm (**c-f**). Corresponds with data in Fig. 2 and 3.

**Extended Data Figure 9.**
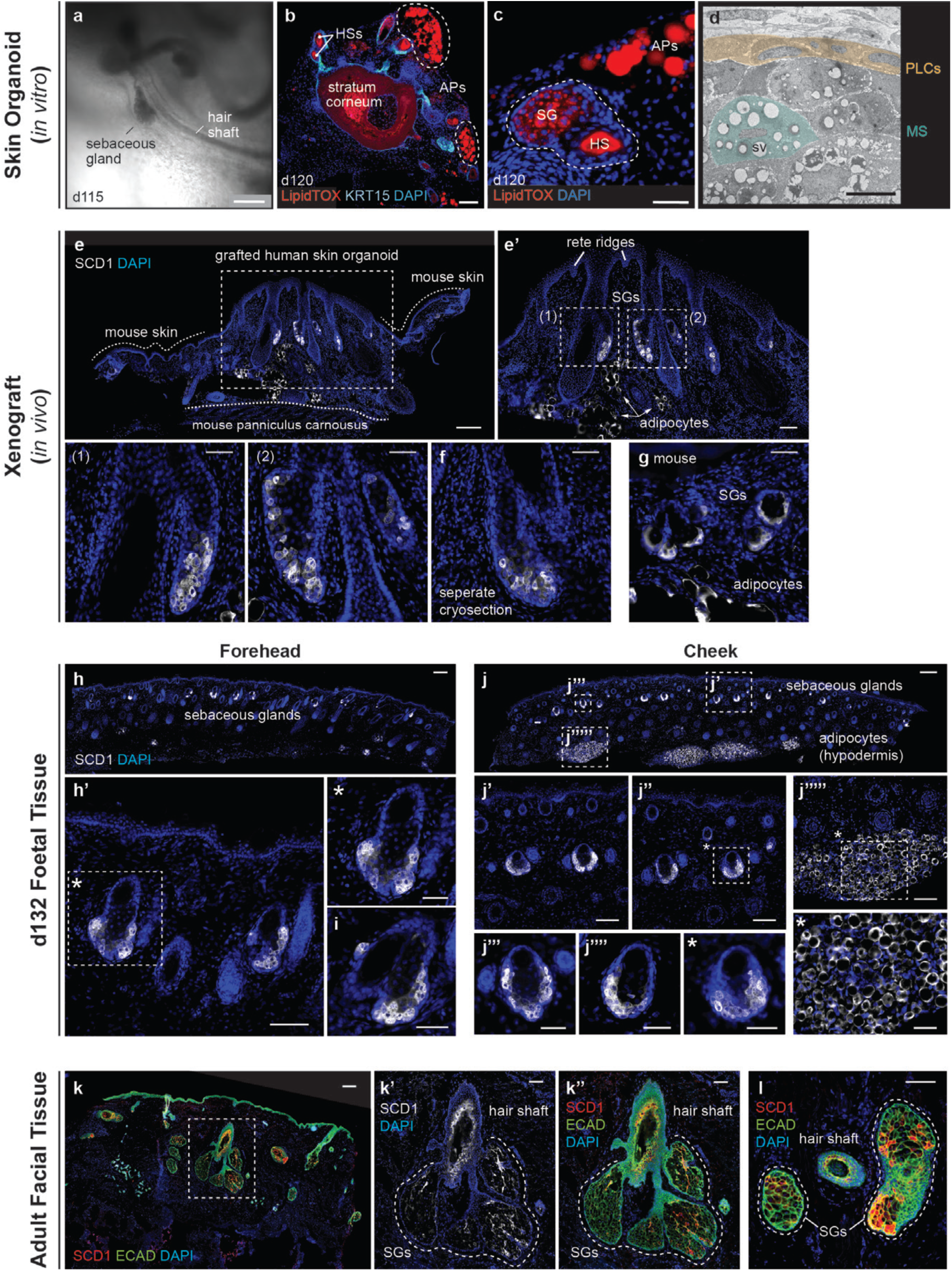
Xenografted WA25 skin organoid HFs have sebaceous glands comparable to second-trimester (day-132) foetal and adult facial hair. **a,** Brightfield image of two day-115 (*in vitro*) WA25 skin organoid HFs with visible hair shafts and sebaceous glands. **b,** LipidTOX staining of day-120 revealing lipid-rich cells, such as sebocytes and adipocytes (APs). KRT15 immunostaining labels outer-root sheath follicle cells. **c,** High magnification image of an adjacent cryosection for the specimen in panel (b). The dashed line highlights a cross-sectioned hair shaft (HS) and sebaceous gland (SG). **d,** TEM image of a day-140 WA25 skin organoid sebaceous gland. Peripheral layer cells (PLCs) and a maturing sebocyte (MS) containing sebum vacuoles (SV) have been pseudo-coloured yellow and green, respectively. **e-f,** SCD1-positive sebaceous glands in xenografted day-140 WA25 skin organoids. Xenografts were extracted and fixed in PFA, >49 days after transplantation. Dashed boxes indicate magnified regions presented in following image panels. Dashed lines distinguish between the grafted human skin organoid tissue and the tissues of host mouse (Nu/J). Arrowheads indicate SCD1^+^ adipocytes. **g,** SCD1-positive sebaceous glands in the NU/J mouse skin adjacent to xenografts. Note that the size of mouse sebaceous glands is smaller than those of human that the origin of sebaceous glands is distinguishable within the extracted xenograft samples. **h-i,** SCD1-positive sebaceous glands in day-132 human foetal forehead skin. **j,** SCD1-positive (j-j’’’’) sebaceous glands and (j’’’’’) adipocytes in day-132 human foetal cheek skin. Note that the adipocytes are prominently abundant in foetal cheek tissue compared to forehead tissue (h vs. j). **k-l,** SCD1-positive sebaceous glands in adult human facial skin. Epithelium is visualized by ECAD immunostaining. Dashed line outlines sebaceous glands of a HF. Scale bars, 250 µm (**a**, **h, j, k**), 100 µm (**b**, **e**, **e’**, **h’, j’, j’’**, **j’’’’’**, **k’**-**l**), 50 μm (**c**, **e’(1)**-**f**, **g**, **h’\***, **i**, **j’’’**, **j’’’’**, **j’’\***, **j’’’’’\***), 5 µm (**d**). Corresponds with data in Fig. 4.

**Extended Data Figure 10.**
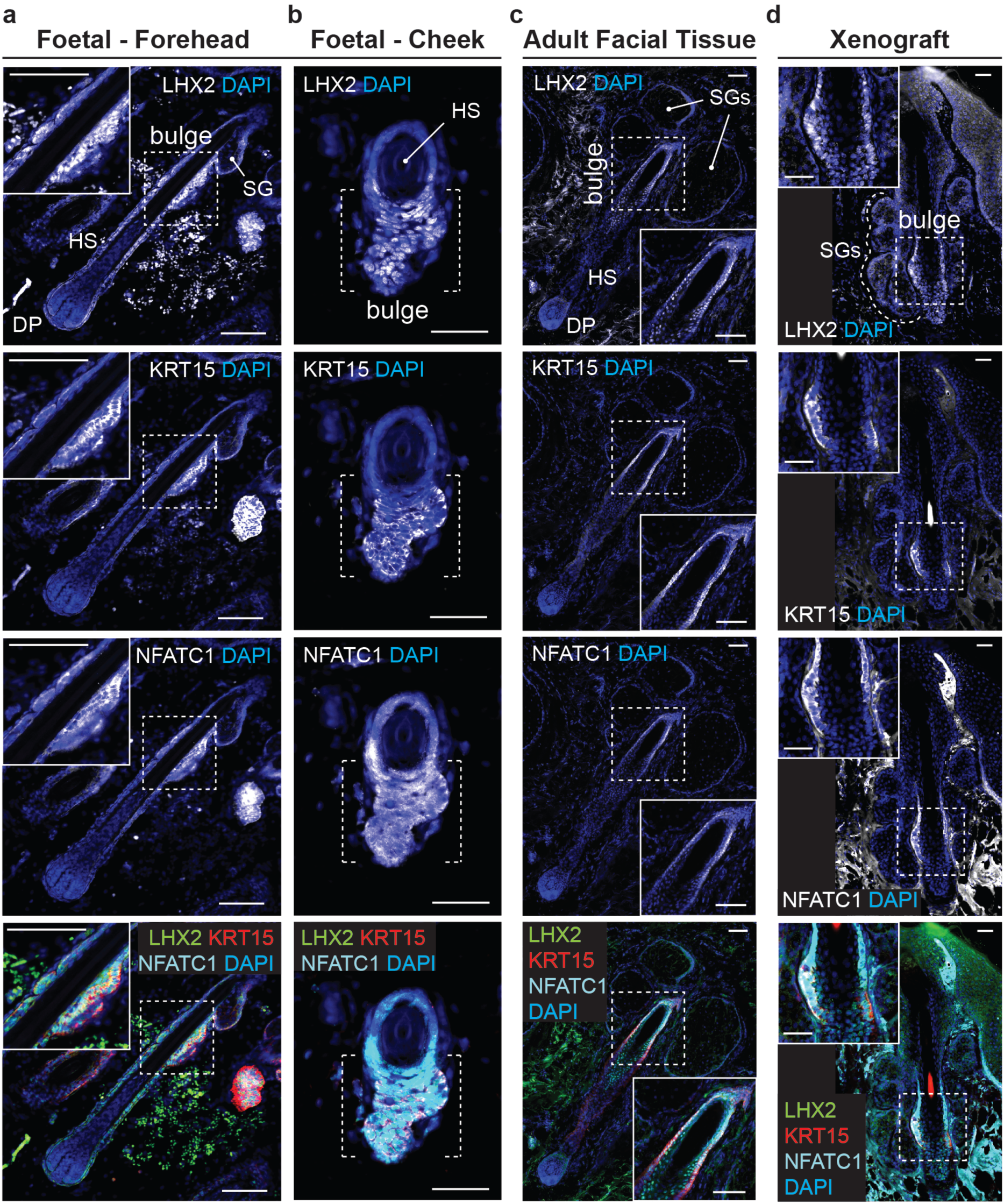
Xenografted WA25 skin organoids have bulge regions comparable to second-trimester (day-132) foetal and adult facial hair. Immunostaining for HF stem cell markers, LHX2, KRT15, and NFATC1, in the HF bulge region of (**a, b**) day-132 human foetal skin from two facial locations (Forehead and Cheek), (**c**) adult facial skin, and (**d**) xenografted skin organoid tissue. **NOTE:** In both foetal and xenograft HFs, NFATC1 expression is predominantly localized to the cytoplasm in bulge cells. By contrast, adult facial HFs have bulge regions have cells with nuclear-localized NFATC1—reminiscent of previous reports showing nuclear-localized NFATC1 in mouse bulge stem cells^4,38^. Dashed boxes indicate magnified regions shown on a corner of each image panel (a, c, d). Dashed brackets show bulge region (b). Abbr: hair shaft (HS); dermal papilla (DP); sebaceous gland (SG). Scale bars, 100 µm (**a**, **c**), 50 µm (**b**, **d**). Corresponds with data in Fig. 4.

