## Supplementary Information for "Hair-bearing human skin generated entirely from pluripotent stem cells"

**SUPPLEMENTARY NOTE**

**Supplementary Note 1 | Extended discussion of methodology development and troubleshooting**

For skin organoid induction, we made substantial modifications to our previously published inner ear organoid protocol[1]. In the Koehler Lab (Indiana University School of Medicine), we primarily used two different human pluripotent stem cell (PSC) lines—WA25 embryonic stem cells (ESCs) and *Desmoplakin-GFP* (*DSP-GFP*) induced PSC (iPSCs)—to optimize the protocol. Then, the optimized protocol was validated in the Heller Lab (Stanford University) using a different cell line, the WA01 (H1) hESC line. We aimed to establish a protocol that was reliable and applicable to multiple PSC lines. Due to the 3D format and the long-duration of the cultures, extensive quantification of cell aggregate morphology and cellular composition was not practical for every treatment regime tested. Therefore, we often used qualitative metrics (e.g. cell aggregate appearance) to discontinue culture conditions at early developmental timepoints, which we judged to be less likely to produce hair-bearing skin organoids. For conditions cultured over 70 days, we have provided the percentage of organoids that displayed hair follicles as a key metric of successful skin organoid generation (see **Supplementary Table 1a**). Below is a brief summary of our optimization approach:

***Basal medium optimization:*** We first sought to substitute an off-the-shelf medium for the Chemically Defined Medium (CDM) used in our inner ear organoid induction protocol. Because CDM was manually prepared for each experiment, we found that pipetting errors, reagent stability, and batch effects were sources of experimental variability. We substituted out CDM for the commercially available Essential 6 (E6) medium (Invitrogen/Gibco) because it is stable, fully defined, and convenient for use. Moreover, E6 medium has been shown to promote highly efficient ectoderm differentiation from PSCs[1]. After switching to E6 medium, however, we noted subtle differences in organoid morphology. Specifically, neural epithelia were more prevalent. Thus, we modified the small molecule and protein treatments for proper skin organoid induction.

***Treatment regime optimization:*** For efficient surface ectoderm formation, the proper concentration and timing of bone morphogenic protein 4 (hereafter, BMP) is critical. In our hands, each PSC line has a different BMP concentration requirement for surface ectoderm induction. This phenomenon is potential due to varying levels of endogenous BMP between cell lines. Therefore, to optimize surface ectoderm formation, we focused on BMP concentration and timing during the first 0 - 4 days of differentiation. We then re-optimized the timing of LDN and FGF treatment. Below is a brief summary of these experiments:

> *BMP concentration/timing optimization*: For *DSP-GFP* cell line, 2.5 – 5 ng/ml of BMP treatment was suitable for skin organoid induction; while 10 ng/ml was also suitable, the outcome was similar to 5 ng/ml BMP. For WA25 cell line, 0 – 2.5 ng/ml of BMP treatment was suitable for skin organoid induction, while 5 ng/ml of BMP was excessive. Thus, we selected 2.5 ng/ml as our universal BMP concentration.

For *DSP-GFP* cells, treatment of BMP was required from the starting day (day 0) of differentiation. BMP treatment beginning on *day 1* partially produced skin organoids, and the treatment on *day 2* or later did not induce skin organoids. For WA25 cell line, BMP treatment timing was more flexible and could be performed any time between *day 0* and *day 1*.

For optimized differentiation of both cell lines, we selected 2.5 ng/ml BMP on the starting day (*day 0*) of differentiation, in addition to the other previously established medium ingredients: E6 basal medium containing 2% Matrigel, 10 μM SB, 4 ng/ml FGF, and 2.5 ng/ml BMP.

*LDN/FGF treatment timing optimization*: To optimize induction of CNCCs, we adjusted the timing of BMP inhibition (LDN treatment) in combination with FGF treatment.

We first examined the timing of LDN/FGF treatment. We found that the required treatment timing varied between *day 3* and *day 4*, depending on the BMP treatment conditions. For WA25 cell line, when BMP was not added, LDN/FGF was required on *day 4* to induce skin organoids. When BMP was added on *day 0*, LDN/FGF was required on *day 3*. In *DSP-GFP* culture, with *day 0* BMP treatment, LDN/FGF treatment on either *day 3* or *day 4* induced skin organoid formation, but *day 3* treatment was more efficient compared to *day 4* treatment (see **Supplementary Table 1b** for results).

We next tested whether FGF treatment would be necessary for efficient skin organoid induction. In WA25 cell line cultures without BMP, LDN treatment on *day 4* induced skin organoids in some cultures, but the efficiency was highly variable from 0 to 66.7% throughout five independent experiments. When LDN was treated in combination with FGF, regardless of treatment timing between *days 4-8*, we consistently observed hair-bearing skin organoids induction: 66.6 ± 11.2% of organoids from five independent experiments ($n$ = 65 organoids). Likewise, in *DSP-GFP* cell line cultures with *day 0* BMP treatment, LDN and FGF co-treatment on *day 3* was more efficient than LDN treatment alone. (see **Supplementary Table 1c** for results).

We concluded the optimal LDN/FGF treatment for both cell lines was 200 nM LDN and 50 ng/ml FGF co-treatment on *day 3* of differentiation.

***Matrigel usage optimization:*** Lastly, we compared Matrigel embedding versus floating culture on *day 12* of differentiation. In our pre-optimized CDM protocol, we embedded individual aggregates in Matrigel (~20 μl per aggregate) droplets as in our inner ear induction protocol. This approach is time consuming and expends large volumes of Matrigel. To overcome these drawbacks, we tested culturing individual aggregates in a floating culture containing 1% Matrigel. We found that 1% Matrigel dissolved in the Organoid Maturation (OM) medium was sufficient to support hair-bearing skin organoid induction. Notably, for this study, we did not test whether skin organoids could be produced in the complete absence of Matrigel at all steps of the protocol.

### SUPPLEMENTARY FIGURE AND LEGEND

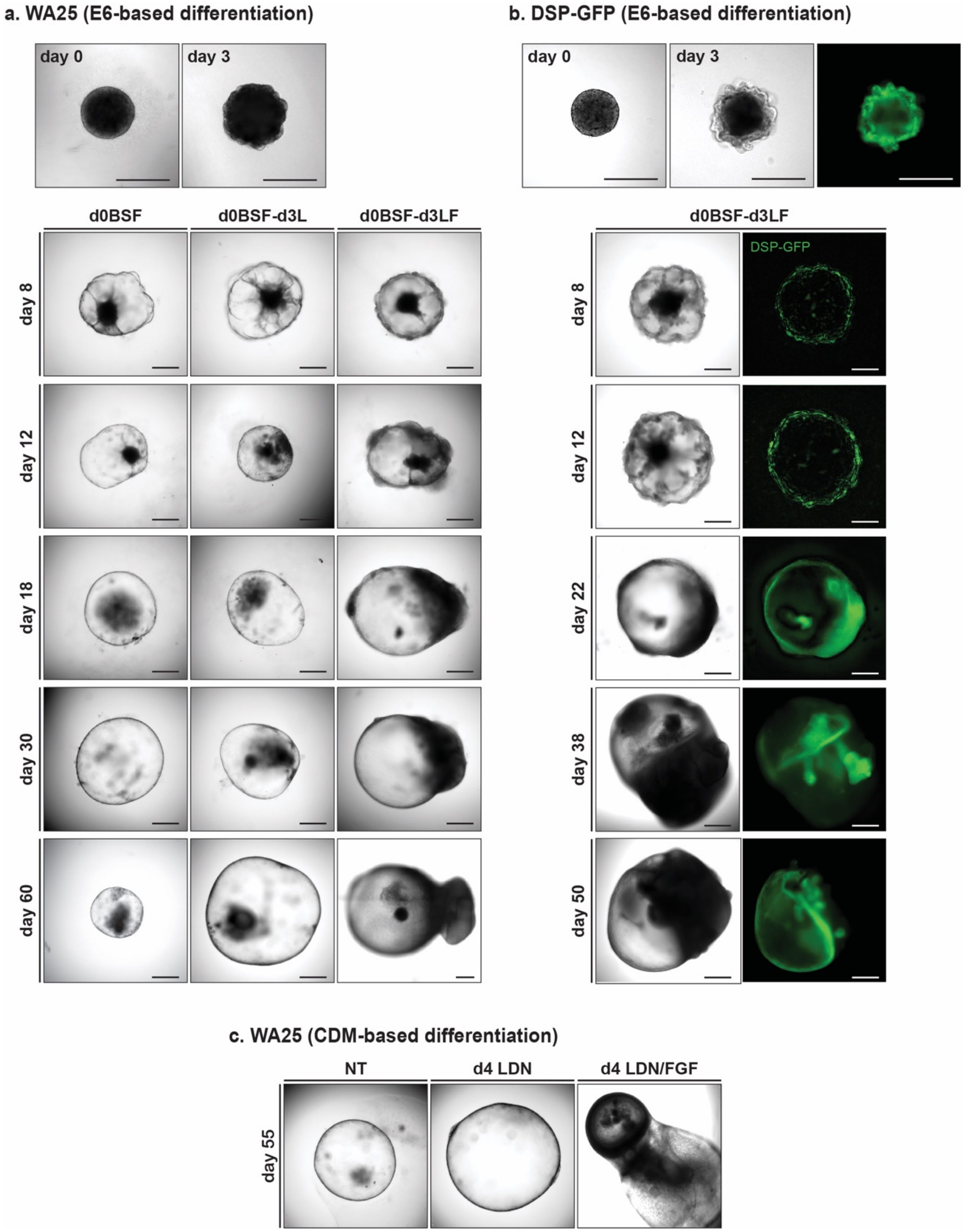


**Supplementary Figure 1: Tracking morphology of skin organoids under different treatment regimes. a,** Representative DIC images of WA25 skin organoid development on differentiation *days 0-60*, cultured in E6-based medium under three different *day 3* treatment conditions – no LDN nor FGF, LDN only, and LDN/FGF. No *day 3* LDN nor FGF treated aggregates maintained cystic organoids for about 30 days of differentiation, but the organoids lost their morphology and shrunk afterwards. Day 3 LDN only treatment supported sphere-like morphology, but was not sufficient enough to produce hair-bearing skin organoids in a consistent manner. Co-treatment of LDN and FGF on day 3 (final optimized treatment condition) was optimal for epithelial stratification and sufficient dermal layer development. Tail-like region containing mesenchymal and neuronal cells on one pole of skin organoids are visible by day 18 of differentiation. **b,** Representative DIC and endogenous GFP fluorescence images of *DSP-GFP* skin organoid development on differentiation *days 0-50*, cultured in E6-based medium with day 3 treatment of LDN/FGF (final optimized treatment regime). $GFP^+$ epithelium is visible on the surface/edge of the sphere-like organoid, and the GFP signal intensifies as the organoids differentiate and mature further. The tail portion of the organoid appears by day 22 of differentiation ($GFP^+$ signal at the tail portion presented in the day 22 image is an autofluorescence). **c,** Representative DIC images of WA25 skin organoid development on differentiation *day 55*, cultured in CDM-based medium under three different *day 4* treatment conditions – no LDN nor FGF, LDN only, and LDN/FGF. Aggregates that were treated with neither LDN nor FGF eventually lost their shape and shrunk. Day 4 LDN treatment maintained cystic organoid morphology, and occasionally induced maturation of skin organoids. Co-treatment of LDN and FGF on day 4 of differentiation was suitable for skin organoids to fully mature in a relatively consistent manner. **Note:** Substitution of CDM- to E6-based differentiation medium improved skin organoid development by reducing the tail portion (non-skin related mesenchymal cells) and increasing the head – skin – portion. **Scale bars, 500 µm (all panels).**

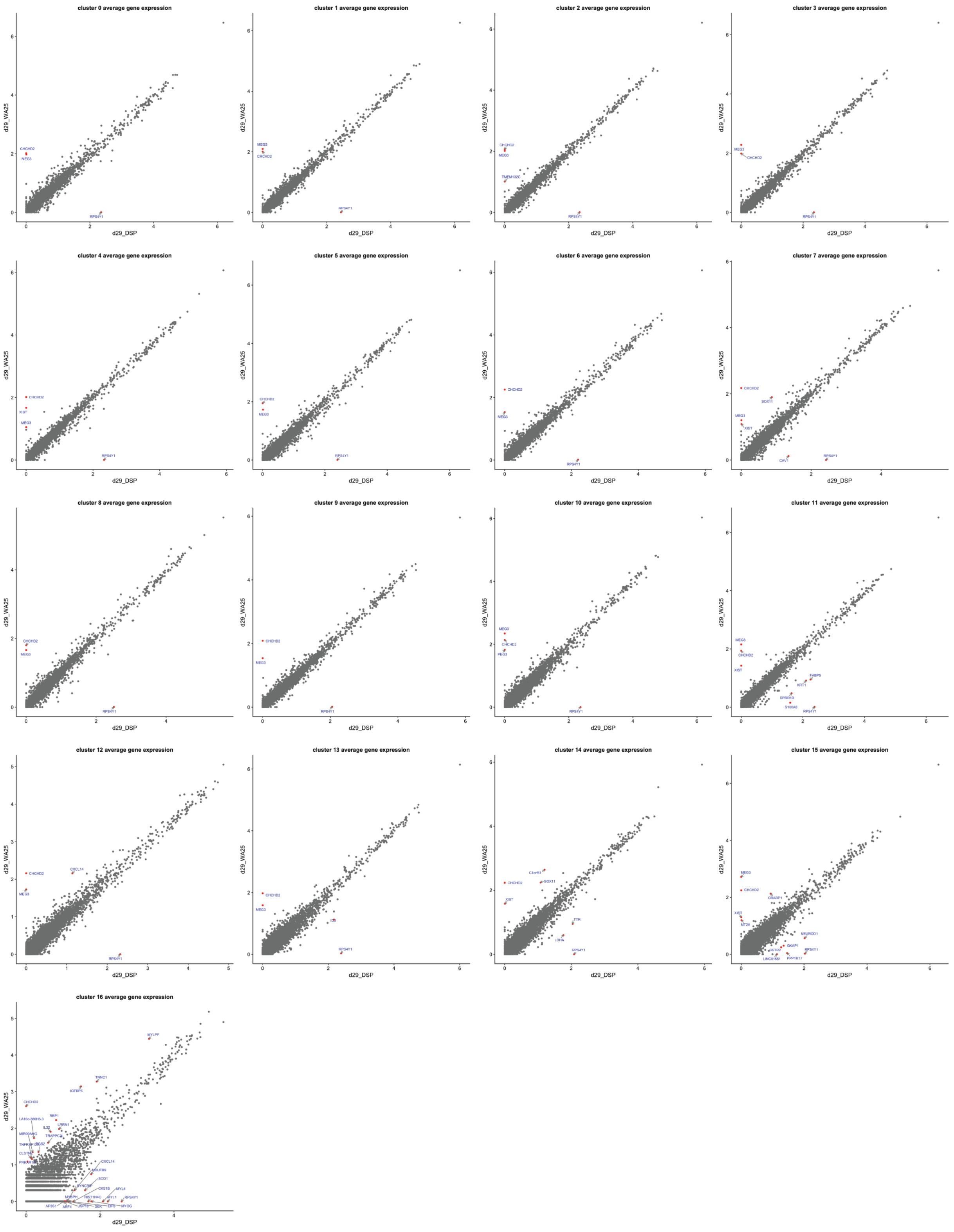


**Supplementary Figure 2: Comparing average gene expression within the same cluster across conditions (day-29 skin organoid datasets).** These graphs show the substantial overlap between genes expressed in cells from WA25 and *DSP-GFP* skin organoids (gray dots). Genes with differential expression—up/down-regulated in WA25 vs. *DSP-GFP*—are indicated in red. WA25 is a female cell line, whereas *DSP-GFP* is a male cell line; therefore, sex-related genes were differentially expressed, such as *XIST* (female) and *RPS4Y1* (male). In WA25, the imprinted genes, *MEG3* and *PEG3*, were upregulated in some cell clusters. Additionally, *CHCHD2* is upregulated in WA25 cells. *CHCHD2* (also known as *MNRR1*) encodes a regulator of mitochondrial metabolism that has been shown to be upregulated in response to cellular stress and/or hypoxic conditions[2]. This gene may be an indication of cellular stress in WA25 cultures; however, additional biological replicates will be needed to confirm this observation. Finally, the lack of correlation between WA25 and *DSP-GFP* gene expression in Cluster 16 (i.e. myocytes) may be due to the relatively low number of WA25-derived myocyte-like cells (3 cells) used for comparison to *DSP-GFP*-derived myocyte-like cells (113 cells).

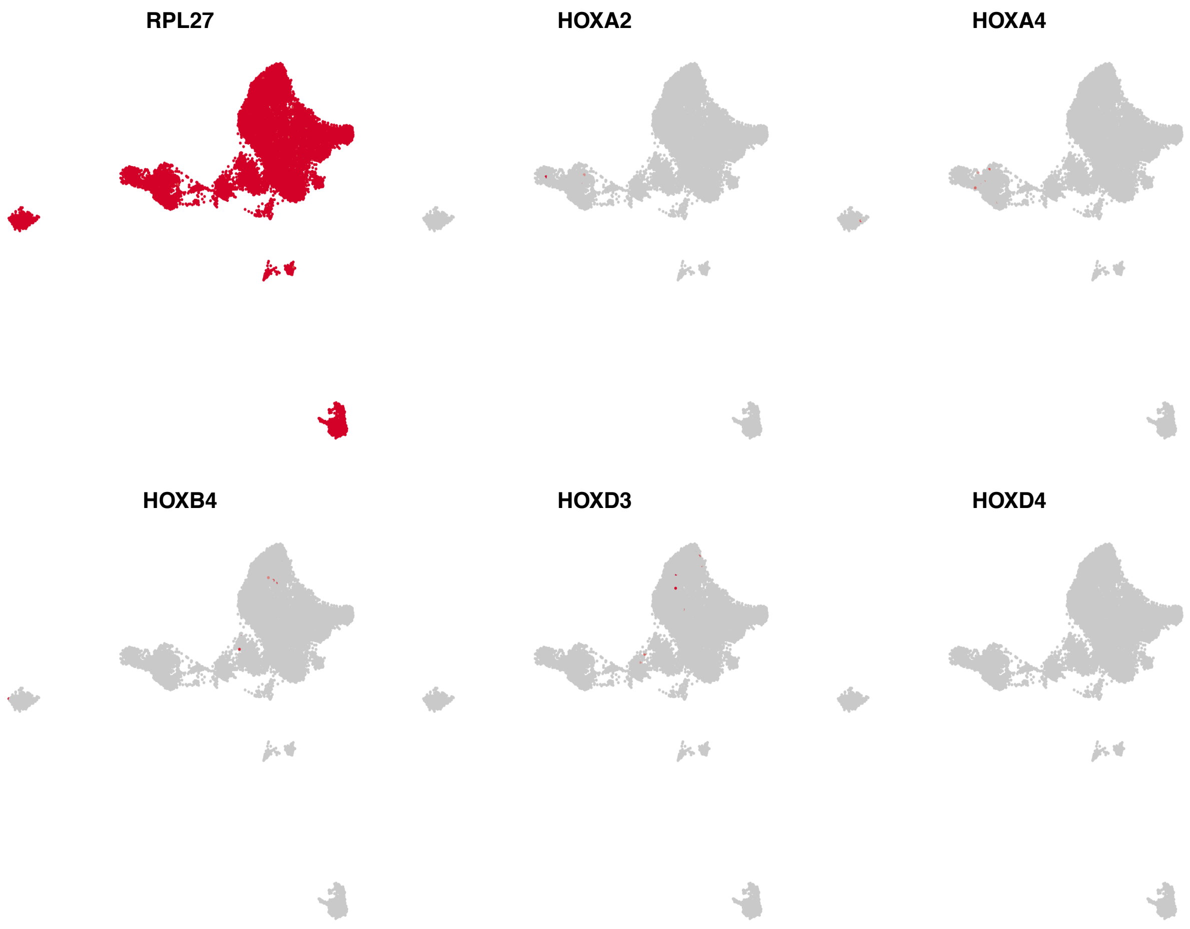


**Supplementary Figure 3: Lack of HOX gene expression in day-29 skin organoids.** There are some cells (<10 cells) within the population that express these HOX genes; however, the vast majority of cells are HOX-negative. The absence of HOX gene expression in a cell subtype does not definitively rule-out expression. There could be significant dropout across all populations for these genes.

### SUPPLEMENTARY TABLES

**Supplementary Table 1 | Comparison of hair follicle formation frequencies between cell lines and treatments.**

**a. Comparison of hair follicle formation frequencies between three different cell lines; WA25 hESCs, DSP-GFP (WTC) hiPSCs, and WA01 hESCs.**

| WA25 hESC | | ***DSP-GFP* hiPSC** | | WA01 hESC | |
|---|---|---|---|---|---|
| **Exp. ID** | **% of HF formation per experiment** | **Exp. ID** | **% of HF formation per experiment** | **Exp. ID** | **% of HF formation per experiment** |
| **#1** | 68.8 | **#1** | 83.3 | **#1** | 70.8 |
| **#2** | 83.3 | **#2** | 95.8 | **-** | - |
| **#3** | 83.3 | **#3** | 79.2 | - | - |
| **#4** | 95.8 | **#4** | 75.0 | - | - |
| **#5** | 88.9 | **#5** | 66.7 | - | - |
| **#6** | 77.8 | **#6** | 84.4 | - | - |
| **#7** | 100 | **#7** | 100 | - | - |
| **#8** | 88.5 | **#8** | 100 | - | - |
| **#9** | 100 | **#9** | 100 | - | - |
| **Average (± SEM)%** | **87.4 (± 3.5)%** | **Average (± SEM)%** | **87.2 (± 4.1)%** | **Average (± SEM)%** | **70.8%** |

**b. Comparison of hair follicle formation frequencies between day 3 vs. day 4 LDN/FGF treatment in WA25 hESCs and *DSP-GFP* hiPSCs.**

| WA25 | | | | | | ***DSP-GFP*** | | | |
|---|---|---|---|---|---|---|---|---|---|
| **d1 BMP + d3 LDN/FGF** | | **d1 BMP + d4 LDN/FGF** | | **d0 BMP + d3 LDN/FGF** | | **d0 BMP + d3 LDN/FGF** | | **d0 BMP + d4 LDN/FGF** | |
| **Exp. ID** | **HF%** | **Exp. ID** | **HF%** | **Exp. ID** | **HF%** | **Exp. ID** | **HF%** | **Exp. ID** | **HF%** |
| **#1** | 85.7 | **#1** | 80.0 | **#1** | 95.8 | **#1** | 100 | **#1** | 91.7 |
| **#2** | 62.5 | **#2** | 100 | - | - | **#2** | 85.7 | **#2** | 83.3 |
| **#3** | 37.5 | **#3** | 87.5 | - | - | - | - | - | - |
| **Average (± SEM)%** | **61.9 (± 13.9)%** | **Average (± SEM)%** | **89.2 (± 5.8)%** | **Average (± SEM)%** | **95.8%** | **Average (± SEM)%** | **92.9 (± 7.1)%** | **Average (± SEM)%** | **87.5 (± 4.2)%** |

**c. Comparison of skin organoid induction efficiency between LDN and LDN/FGF treatment conditions in WA25 hESCs and *DSP-GFP* hiPSCs.**

| WA25 | | | | *DSP-GFP* | | | |
|---|---|---|---|---|---|---|---|
| No BMP + d4 LDN | | No BMP + d4 LDN/FGF | | d0 BMP + d3 LDN | | d0 BMP + d3 LDN/FGF | |
| Exp. ID | HF% | Exp. ID | HF% | Exp. ID | HF% | Exp. ID | HF% |
| **#1** | 66.7 | **#1** | 83.3 | **#1** | 80 | **#1** | 100 |
| **#2** | 8.3 | **#2** | 91.7 | **-** | | **-** | |
| **#3** | 0 | **#3** | 37.5 | **-** | | **-** | |
| **#4** | 0 | **#4** | 78.6 | **-** | | **-** | |
| **#5** | 36.4 | **#5** | 42.1 | **-** | | **-** | |
| **Average (± SEM)%** | **22.3 (± 13)%** | **Average (± SEM)%** | **66.6 (± 11.2)%** | **Average (± SEM)%** | **80%** | **Average (± SEM)%** | **100%** |

### Supplementary Table 2 | Media compositions.

#### a. Pre-aggregation Media

- **For cell collection (10Y)**

| Component | Supplier | Cat. No. | Stock Concentration | Final Concentration | Volume (10 ml) |
|---|---|---|---|---|---|
| Essential 8 Flex Medium | Gibco | A2858501 | - | 100% (v/v) | 10 ml |
| Y27632 in Solution | Stemgent | 04-0012-02 | 10 mM | 10 µM | 10 µl |
| Normocin | Invivogen | Ant-nr-1 | 50 mg/ml | 100 µg/ml | 20 µl |

- **For cell seeding (20Y)**

| Component | Supplier | Cat. No. | Stock Concentration | Final Concentration | Volume (22 ml) |
|---|---|---|---|---|---|
| Essential 8 Flex Medium | Gibco | A2858501 | - | 100% (v/v) | 22 ml |
| Y27632 in Solution | Stemgent | 04-0012-02 | 10 mM | 20 µM | 44 µl |
| Normocin | Invivogen | Ant-nr-1 | 50 mg/ml | 100 µg/ml | 44 µl |

**b. Skin Organoid Induction Medium**

- **Chemically defined medium (CDM)**

| Component | Supplier | Cat. No. | Stock Concentration | Final Concentration | Volume (100 ml) |
|---|---|---|---|---|---|
| Ham's F-12 Nutrient Mix, GlutaMAX™ Supplement | Gibco | 31765-035 | - | 49% (v/v) | 49 ml |
| IMDM, GlutaMAX™ Supplement | Gibco | 31980-030 | - | 49% (v/v) | 49 ml |
| Chemically Defined Lipid Concentrate | Invitrogen | 11905-031 | 100X | 1X | 1 ml |
| Bovine Serum Albumin (BSA) | Sigma | A1470 | - | 5 mg/ml | 0.5 g |
| Insulin Solution Human | Sigma | I9278 | 10 mg/ml | 7 µg/ml | 70 µl |
| Transferrin Human | Sigma | T8158 | 20 mg/ml | 15 µg/ml | 75 µl |
| 1-Thioglycerol | Sigma | M6145 | 11.5 M | 450 µM | 4 µl |
| Normocin | Invivogen | Ant-nr-1 | 50 mg/ml | 100 µg/ml | 200 µl |

*Note: For day 0 differentiation, 600 µl Matrigel (2%) is added to 30 ml of CDM, a volume of which is needed for transferring all aggregates from 2 of 96-well U-bottom plates (192 aggregates) to 2 of new 96-well U-bottom plates. For treatments on day 3 and fresh medium additions on day 8, CDM without Matrigel is used from the stock (remaining 70 ml after use on day 0) made ahead on day 0. Use the medium within 10 days – make fresh CDM prior to initiating experiments.*

- **E6-based differentiation medium**

| Component | Supplier | Cat. No. | Stock Concentration | Final Concentration | Volume (30 ml) |
|---|---|---|---|---|---|
| Essential 6 Medium | Gibco | A1516401 | - | 98% (v/v) | 29.4 ml |
| Matrigel | Corning | 354230 | - | 2% (v/v) | 600 µl |
| SB431542 (SB) | Stemgent | 04-0010-05 | 10 mM | 10 µM | 30 µl |
| Recombinant Human FGF-basic (FGF) | PeproTech | 100-18B | 200 µg/ml | 4 ng/ml | 0.6 µl |
| Recombinant Human BMP-4 (BMP) | PeproTech | 120-05 | 100 µg/ml | 2.5 ng/ml | 0.75 µl |
| Normocin | Invivogen | Ant-nr-1 | 50 mg/ml | 100 µg/ml | 60 µl |

*Note: For treatments on day 3 and half-medium changes on day 8, E6 medium containing only Normocin (antibiotics) was freshly prepared and used. Be aware of expiration dates of each small molecule and protein – use within 6 months of receiving and/or reconstituted date.*

**c. Skin Organoid Maturation Medium (OMM)**

| Component | Supplier | Cat. No. | Stock Concentration | Final Concentration | Volume (50 ml) |
|---|---|---|---|---|---|
| Advanced DMEM/F12 | Gibco | 12634010 | - | 49% (v/v) | 24.5 ml |
| Neurobasal Medium | Gibco | 21103049 | - | 49% (v/v) | 24.5 ml |
| GlutaMAX™ Supplement | Gibco | 35050061 | 100X | 1X | 500 µl |
| B-27 Supplement, Minus Vitamin A | Gibco | 12587010 | 50X | 0.5X | 500 µl |
| N2 Supplement | Gibco | 17502048 | 100X | 0.5X | 250 µl |
| 2-Mercaptoethanol | Gibco | 21985023 | 55 mM | 0.1 mM | 91 µl |
| Normocin | Invivogen | Ant-nr-1 | 50 mg/ml | 100 µg/ml | 100 µl |

*Note: For day 12 (transition of each aggregate into 24-well low-attachment plates in 500 µl of OMM per well and start consistent agitation of the culture for better perfusion of medium) and day 15(half-medium changes), use OMM containing 1% Matrigel. From day 18 to 30, perform half-medium changes every three days, and afterwards, every other day with full-medium change once a week. Make fresh OMM and use within a week.*

**Supplementary Table 3 | Antibodies list. Related to Experimental Procedures.**

| Protein | Host | Vendor | Catalog No. | Dilution | Markers for: |
|---|---|---|---|---|---|
| Aggrecan (ACAN) | Rabbit | Invitrogen | PA1-1745 | 1:100 | Chondrocytes |
| βIII-Tubulin (TUJ1) | Mouse | BioLegend | 801202 | 1:100 | Neurons |
| CD34 | Rat | BD Biosciences | 553731 | 1:50 | Dermal fibroblasts |
| CD49f | Rat | Invitrogen | 12-0495-81 | 1:50 | Epidermal layer |
| Collagen 2A1 (COL2A1) | Mouse | Invitrogen | MA1-37493 | 1:10 | Chondrocytes |
| Cytokeratin 5 (KRT5) | Rabbit | Invitrogen | MA5-14473 | 1:50 | Basal layer of skin |
| Cytokeratin 5 (KRT5) | Mouse | Invitrogen | MA5-12596 | 1:50 | Basal layer of skin |
| Cytokeratin 10 (KRT10) | Mouse | Santa Cruz | sc-23877 | 1:50 | Spinous layer of skin |
| Cytokeratin 15 (KRT15) | Mouse | Santa Cruz | sc-47697 | 1:50 | Periderm and basal layers of skin |
| Cytokeratin 17 (KRT17) | Mouse | Santa Cruz | sc-393091 | 1:50 | Periderm, epidermis & outer root sheath |
| Cytokeratin 20 (KRT20) | Rabbit | Cell Signaling | 13063S | 1:100 | Markel cells |
| E-cadherin (ECAD) | Mouse | BD Biosciences | 610181 | 1:50 | Epithelium |
| EDAR | Goat | R&D Systems | AF745-SP | 1:50 | Hair placodes |
| Islet1 (ISL1) | Mouse | DSHB | 39.4D5 | 1:5 | Merkel progenitor cells & sensory neurons |
| Ki67 | Mouse | BD Biosciences | 550609 | 1:100 | Proliferating cells |
| LHX2 | Rabbit | Millipore | ABE1402 | 1:750 | Hair placodes & bulge |
| Loricrin | Rabbit | Abcam | ab85679 | 1:50 | Granular layer of skin |
| Nephronectin (NPNT) | Mouse | R&D Systems | MAB8025 | 1:50 | Arrector pili muscle |
| Neurofilament-heavy chain (NEFH) | Mouse | Cell Signaling | 2836S | 1:100 | Sensory neurons |
| NFATC1 | Mouse | Invitrogen | MA3-024 | 1:50 | Hair follicle bulge |
| P-cadherin (PCAD) | Mouse | Invitrogen | 32-4000 | 1:50 | Hair placodes |
| p75NTR (P75) | Rabbit | Cell Signaling | 8238 | 1:50 | Neural crest cells & dermal condensates |

| | | | | | |
|---|---|---|---|---|---|
| PDGFRα (D13C6) | Rabbit | Cell Signaling | 5241 | 1:50 | Dermal fibroblasts |
| PMEL | Mouse | Novus | NBP2-29407 | 1:100 | Melanocytes |
| S100β | Rabbit | Abcam | ab52642 | 1:75 | Schwann cells |
| SCD1 | Rabbit | Cell Signaling | 2794 | 1:100 | Sebaceous glands |
| SOX2 | Mouse | BD Biosciences | 561469 | 1:100 | Dermal papilla/condensates, Merkel cells, Melanocytes |
| SOX10 | Mouse | Invitrogen | 14-5925-80 | 1:50 | Neural crest cells |
| TFAP2α | Mouse | DSHB | 3B5 | 1:5 | Epithelial & cranial neural crest cells |

*Note: All antibodies used for IHC were previously validated in IHC experiments. Citations can be found on the manufacture's website.*

### SUPPLEMENTARY VIDEO LEDGENDS

**Supplementary Video 1 | Day 85 WA25 skin organoids with nascent hair follicles.** Wholemount skin organoid immunostained with antibodies for KRT5 (red) and SOX2 (cyan). The sample was cleared using the ScaleS approach and imaged on an Olympus FV1000 confocal microscope. The dense clusters of $SOX2^+$ cells represent the dermal condensate and dermal papilla cells. The $SOX2^+$ cells embedded in the outer root sheath of each follicle are presumed to be Merkel cells, while the extra-epithelial $SOX2^+$ cells are presumed to be melanoblasts.

**Supplementary Video 2 | Day 110 WA25 skin organoid with dermal papilla cells and neurites.** Wholemount skin organoid immunostained with antibodies for KRT5 (red), SOX2 (cyan), and TUJ1 (green). Image segmentation using the Imaris "spots" module to estimate that the dermal papilla cell (DPC) clusters contained approximately 250, 335, and 522 cells.

**Supplementary Video 3 | *DSP-GFP* skin organoid with $PMEL^+$ melanocytes.** Wholemount skin organoid immunostained with an antibody for PMEL (red).

**Supplementary Video 4 | Neural network in nascent day 85 WA25 skin organoid.** Wholemount skin organoid immunostained with an antibody for TUJ1 (red). Hoechst staining is displayed as cyan.

**Supplementary Video 5 | Mechanosensory complexes in a day 140 WA25 skin organoids.** Wholemount skin organoid immunostained with antibodies for KRT17 (red) and TUJ1 (green).

**Supplementary Video 6 | NPNT expression in an organoid HF bulge region.** Wholemount skin organoid immunostained with antibodies for KRT17 (red), KRT20 (green), and NPNT (white). Note the high cell density of outer root sheath cells in the bulge region compared to non-bulge region (upper part of the frame).
